# Targeted mining of plastic-associated metagenomes uncovers a novel thermostable PETase expanding scaffold space for engineering

**DOI:** 10.64898/2026.07.10.737215

**Authors:** Konstantinos Rigkos, Dimitra Bezantakou, Kyriakos Antoniadis, Io Antonopoulou, Dimitra Zarafeta, Georgios Skretas

## Abstract

Enzymatic depolymerization of polyethylene terephthalate (PET) has advanced rapidly, alongside a growing volume of publicly available metagenomic data from microbial communities under sustained selective pressure from plastic exposure. Reasoning that such environments may harbor underexplored polyester-active enzymes, we developed a targeted mining workflow that screens exclusively plastic-associated datasets through multi-step bioinformatic filtering—integrating catalytic-motif screening, disulfide-topology validation, structural-similarity scoring, and phylogenetic profiling—to recover high-confidence PETase candidates. Applied to 277 plastic-associated metagenomes, the pipeline yielded 21 non-redundant candidates, several of which combine the Type I catalytic motif (GHSMGGGG) with Type II-like extended loops and secondary disulfide bonds. Two candidates were experimentally confirmed as PET hydrolases; the more active, PET-KR1, is a thermostable enzyme (*Tm* = 66.5 °C) that depolymerizes PET across a broad temperature range, with markedly higher productivity on powdered than on film substrate. PET-KR1 achieved optimal depolymerization at 50 °C, yet at 60–65 °C, where total yields declined, the product pool was more strongly enriched in the terminal monomer TPA, suggesting that thermostability and substrate accessibility are the primary targets for further engineering. Molecular dynamics simulations revealed a conserved hydrophobic binding network around the catalytic serine, consistent with established PETase substrate-recognition modes, and rational disulfide engineering raised the melting temperature by 3.5 °C, confirming amenability to further optimization. Overall, PET-KR1 expands the scaffold space available for PETase engineering, while the discovery workflow, built entirely on publicly available tools and open-access data, provides a reproducible strategy for metagenomic mining of novel PET-degrading enzymes toward biocatalytic PET recycling.

## Introduction

Global plastic production has risen from 2 million tons in 1950 to over 430 million tons in 2024 ^1^. It is estimated that 60% of the total produced plastics have been discarded in landfills and the environment with 4.8 to 12.7 million tons entering the oceans each year ^2,3^. Postconsumer plastic waste is mainly managed by landfill disposal (∼25%), incineration (∼40%), and recycling (<30%) ^2,4^. Incineration decreases waste volume but releases harmful emissions, making it less favorable despite energy recovery ^5^. Growing environmental awareness has driven the demand for improved recycling technologies and policies to reduce plastic use and enhance recyclability ^6–8^.

Plastics are currently recycled mainly through mechanical or chemical routes ^9^. While mechanical recycling can reduce environmental impacts, it is significantly limited by the complexity of plastic compositions, inefficient sorting processes, and the reduced quality of the recycled products ^9,10^. Chemical recycling, on the other hand, holds promise for advancing a sustainable circular economy ^11^. However, its implementation is currently constrained by high energy demands and the toxicity of chemical catalysts, which translate into elevated operational costs and environmental footprint ^11,12^. Addressing these challenges requires innovative approaches to increase recycling rates and simultaneously establish techno-economic feasibility and sustainability.

Biocatalytic recycling, which leverages microorganisms or enzymes to depolymerize plastics, presents an eco-friendly alternative to conventional recycling methods ^13^. In particular, enzyme-catalyzed depolymerization has shown significant promise, with 495 documented plastic-active enzyme activities cataloged in PAZy: The Plastics-Active Enzymes Database ^14^. Among these, 320 enzymes are identified as polyethylene terephthalate (PET) hydrolases (PETases), with others displaying specific activity against polyurethane (PUR) and polyamide (PA) ^14^. Notably, other types of plastics containing a carbon-carbon backbone remain recalcitrant to biocatalytic cleavage ^15^. The focus on PET-degrading enzymes is driven by the global prevalence of PET, one of the most widely used types of plastic ^16^. The structure of PET comprising repeating units of terephthalic acid (TPA) and ethylene glycol (EG) linked by ester bonds—gives it excellent properties for textile and packaging industries, such as high mechanical, thermal and chemical stability ^17–19^. However, these same properties render PET a persistent pollutant that is challenging to degrade and recycle ^20^. PET waste and pollution has been detected in various environmental compartments, including soils, sediments, groundwater, and marine ecosystems, in both macroplastic and microplastic forms ^21^. Moreover, it has been shown to bioaccumulate within living organisms through the food chain, leading to harmful metabolic effects ^21,22^. Therefore, the discovery and engineering of PET-degrading enzymes are crucial for the development of biocatalytic recycling technologies aimed at mitigating the challenges posed by plastic waste.

Benchmark engineered PETases, such as LCC-ICCG and Kubu-PM12, demonstrate industrial viability, achieving >90% amorphous PET conversion at 70 °C ^23,24^. The company Carbios has advanced enzymatic recycling to near-commercial scale, with its delayed Longlaville plant (50 kt/year capacity) targeting production start by H1 2028 and a first licensing agreement for a plant in China announced in 2026, underscoring the commercial momentum behind enzymatic PET depolymerization ^25,26^. These key advancements highlight the continuous maturation of enzyme technology towards the more sustainable handling of plastic waste. Yet, further expansion of the repertoire of the available PET-degrading enzymes diversity remains essential for understanding and optimizing performance across diverse waste streams, crystallinity levels, and process conditions. A key determinant of process performance is thermostability. Efficient enzymatic PET depolymerization typically requires sustained activity near the glass-transition temperature of amorphous PET (Tg ≈ 70 °C), where increased polymer chain mobility enhances enzyme access to ester bonds — making thermostability a critical criterion in both the discovery and engineering of new PETase candidates ^27^.

Over the past decade, PETase discovery has evolved from targeted culture isolation to more sophisticated and high-throughput computational and metagenomic strategies. The archetypal PET-degrading enzyme IsPETase, isolated from *Ideonella sakaiensis* grown on PET bottles ^28^, and the thermostable PETase LCC derived from leaf-branch compost metagenomes ^29^ established the field, yet their performance limitations spurred diverse discovery paradigms. Recent advances include sequence clustering of 1,894 candidates yielding industrial enzymes like Kubu-PM12 ^24^, machine learning-guided screening of 212 homologs identifying 91 PETases including novel low-pH actives ^30^, structure-based deep learning discovering 14 PETases including thermostable enzymes like KbPETase ^31^, marine metagenome mining revealing novel clades ^32,33^, and ancestral reconstruction mapping evolutionary pathways ^34^.

While these approaches have greatly expanded the PETase sequence space, they have primarily drawn on general protein databases, environmental metagenomes, or reconstructed ancestors, without yet fully leveraging (meta)genomic datasets from environments exposed to plastics. Anthropogenic plastic debris represents a persistent and chemically homogeneous carbon source in natural environments. Microbial communities colonizing plastic surfaces therefore experience sustained exposure to polyester polymers and associated additives. Given the relatively recent and widespread introduction of plastic materials into the environment, we reasoned that prolonged exposure to synthetic polyesters could impose a selective pressure favoring microorganisms that harbor enzymes capable of acting on these substrates ^35,36^. We therefore hypothesized that plastic-associated metagenomes represent a promising but underexplored reservoir for polyester-active hydrolases, and that targeted mining of these datasets might recover PETase candidates with features distinct from those identified in general sequence databases or non-plastic environments.

To address this gap, in the present work, we have exclusively targeted plastic-associated metagenomic datasets and have analyzed them bioinformatically using *ProteoSeeker* ^37^, our open source metagenomic protein discovery platform. Through a multi-step screening strategy—incorporating PETase-specific sequence motifs, structural similarity to benchmark PETases, molecular docking, and phylogenetic profiling—we identified 21 high-confidence candidates.

Experimental validation revealed two novel PET-hydrolyzing enzymes, including PET-KR1, a thermostable PETase (*Tm* = 66.5 °C). PET-KR1 additionally combines a Type I catalytic motif with Type II-like structural features, illustrating the architectural diversity recoverable through this approach. Biochemical characterization, molecular simulations, and rational engineering were subsequently employed to investigate its catalytic properties and potential as an engineering scaffold. Beyond the discovery of PET-KR1, our findings establish plastic-associated metagenomes as a tractable and still largely untapped reservoir for PET-degrading enzymes and provide a reproducible workflow for mining them.

## Materials and Methods

All chemical reagents used in this study were purchased from AppliChem and all molecular biology related products were obtained from New England Biolabs unless stated otherwise. Amorphous PET film (aPET; Goodfellow Ltd., GF25214475) was purchased from Merck.

### Bioinformatic Screening

Metagenomic datasets associated with plastic environments were retrieved from the European Nucleotide Archive (ENA) ^38^ using the following query in advanced search for Raw reads as Data type: (sample_description=“plastic” OR sample_title=“plastic”) AND library_strategy=“WGS” AND library_source=“METAGENOMIC”. The ENA search returned 334 RUN accession codes (search performed in November 2024), which were subsequently curated to exclude entries with non-plastic-associated sample descriptions or otherwise irrelevant metadata, resulting in a final set of 277 metagenomes for downstream analysis.

Raw next-generation sequencing reads were analyzed using *ProteoSeeker* (v1.0) ^37^ in both seek and taxonomy modes. Seek mode performed targeted protein domain screening to identify sequences matching user-specified Pfam ^39^ families from assembled contigs, while taxonomy mode generated comprehensive taxonomic classification across all predicted proteins.

*ProteoSeeker* performs automated quality control, contig assembly via MEGAHIT ^40^, gene prediction using FragGeneScanRs ^41^, and protein family screening via HMMER ^42^ against user-specified protein domains. In November 2024, screening targeted four Pfam accessions corresponding to protein families enriched in characterized PETases: Cutinase (PF01083.26), Chlorophyllase (PF07224.15), Chlorophyllase2 (PF12740.11), and Dienelactone hydrolase family (PF01738.22), reflecting the absence of dedicated PETase domains at that time. These proxy families were selected based on their representation in biochemically characterized PET hydrolases (IsPETase, LCC, PHL7) ^43^.

This analysis identified >77 million putative proteins across all 277 metagenomes. Pooled seek (Pfam-targeted) and taxonomy (domain-agnostic) outputs were filtered to identify putative PET-degrading enzymes using the following criteria: (i) presence of conserved sequence motifs (Gly-His-Ser-Met-Gly-Gly-Gly-Gly and Gly-Trp-Ser-Met-Gly-Gly-Gly-Gly), historically associated with Type I-like and Type II-like PETases ^13^, respectively, and (ii) length between 250–350 amino acids, consistent with the size range of characterized PETases. These filters yielded 54 candidates, from which N-terminal signal peptides were predicted and trimmed using Phobius ^44^.

Candidate sequences were further evaluated for the presence of a C-terminal disulfide bond, a common structural feature of characterized PETases, known to contribute to enzyme stability and catalytic efficiency. Specifically, sequences were first screened for the presence of at least two cysteine residues in the C-terminal tail region. Selected candidates were then aligned against a reference set of known PET hydrolases (IsPETase, LCC, and TfCut2) using MAFFT Multiple sequence alignment (MSA) ^45^ to verify cysteine positioning and validate putative disulfide bond topology. Additionally, the presence of Ser-His-Asp catalytic triad and other features historically associated with Type II-like PETases, such as extended loops and a putative secondary disulfide, were evaluated. This stringent filtering resulted in 30 putative PETases, which were then clustered using CD-HIT (v4.8.1) ^46^ with a sequence identity threshold of 100%. This approach removed exact sequence duplicates, yielding a final set of 21 unique putative PETase sequences.

As a final structural validation, predicted structural models of all 21 unique sequences were generated using ColabFold (AlphaFold 2 with MMseqs2) ^47,48^ and superimposed with the crystal structure of LCC (PDB: 4EB0) ^49^ via TM-Align ^50^. All candidates exhibited TM-scores ≥ 0.9 (mean 0.95 ± 0.01), demonstrating highly conserved α/β-hydrolase folds consistent with PETase architecture.

### Sequence Similarity, Phylogenetic Analysis and sequence features

All 21 putative PETases were analyzed by BLASTP ^51^ against the Swiss-Prot and nr databases (2025_12 release). Additionally, 2,575 reference sequences compiled from three landmark PETase discovery studies published in 2025 by Seo *et al*., Norton-Baker *et al*. and Wu *et al*. were included in the analysis ^24,30,31^. Key metrics extracted included percentage identity (%ID), query coverage, and E-values to identify novel PETase candidates. For each query, the top hit was selected based on highest sequence identity where coverage remained high (≥82%), prioritizing functionally relevant homologs with maximal similarity across near-complete protein lengths.

Phylogenetic analysis was conducted by utilizing MAFFT MSA for the 21 putative PETases and 13 characterized reference PETases (IsPETase, LCC, TfCut2, PHL7, and homologs). α/β-hydrolase superfamily references—including cutinase (CUTI1_FUSVN), lipase (MDLA_PENCA), and acetyl xylan esterase (AXE2_GEOSE)—contextualized PETase placement, with classical dienelactone hydrolases (DLH variants) serving as stable outgroups. Maximum-likelihood phylogenetic analysis was performed in IQ-TREE 3.0.1 ^52^ (-m TEST -bb 1000 -nt AUTO), with ModelFinder ^53^ running the optimal LG+G4 substitution model and 1000 ultrafast bootstrap replicates for branch support. Trees were visualized in iTOL v6 ^54^, revealing PETase-linked clades rooted against stable outgroups, with candidates distributed across established Type I/II radiations and putative intermediate lineages.

To identify key functional PETase motifs at the sequence level, MSA was performed on the 21 putative PETases plus 15 reference PETases (IsPETase, LCC, Mipa-P, Kubu-P, PHL7, etc.) using MAFFT. The resulting alignment was visualized using the ESPript 3.0 server ^55^ to annotate critical structural and functional characteristics, including the conserved catalytic triad (Ser-His-Asp) essential for ester hydrolysis, distinctive Type I/II motifs (GHSMGGGG/GWSMGGGG) that define PETase structural subclasses, C-terminal disulfide cysteines important for stability, and characteristic extended loops.

### Recombinant Expression and Purification of PETase Candidates

Genes encoding PETcand1 (PET-KR1), PETcand9, PETcand15, and PETcand16 were codon-optimized for *Escherichia coli* expression and synthesized commercially in the pET-29b(+) vector (Twist Bioscience), generating pET-PETcand(1-16) constructs. The inserts comprised the signal-peptide-free (mature) enzyme sequences cloned between the *Nde*I and *Xho*I restriction sites, resulting in proteins bearing a C-terminal 6XHis tag for affinity purification.

*E. coli* SHuffle® T7 Express chemically competent cells (New England Biolabs) were transformed with the recombinant pET-PET_candX_ plasmids to enable cytosolic disulfide bond formation and recombinant protein production. Cultures were grown in LB medium supplemented with kanamycin (50 μg mL⁻¹) at 37 °C and 220 rpm to an OD₆₀₀ of 0.6–0.8. At this stage protein production was induced by addition of 0.5 mM isopropyl-β-D-thiogalactopyranoside (IPTG), followed by incubation at 18 °C for 16–20 h with shaking at 220 rpm.

Cells were harvested by centrifugation (4,000 × g, 15 min, 4 °C) and resuspended in lysis buffer (25 mM Tris-HCl, 100 mM NaCl, 10 mM imidazole, pH 8.3). Cells were lysed by sonication on ice using a SONICS Vibra-Cell ultrasonic processor at 100% amplitude for 20 cycles (20 s on/20 s off). The resulting lysates were clarified by centrifugation (17,000 × g, 45 min, 4 °C) and applied to an immobilized metal affinity chromatography (IMAC) column packed with Protino Ni-NTA Agarose (MACHEREY-NAGEL). The column was equilibrated with lysis buffer, washed with buffer containing 25 mM Tris-HCl, 100 mM NaCl, 30 mM imidazole (pH 8.3), and bound proteins were eluted with 25 mM Tris-HCl, 100 mM NaCl, 300 mM imidazole (pH 8.3). The PETase content (% w/w) in the purified protein fraction after IMAC purification was determined by SDS-PAGE using Bovine Serum Albumin as protein standard (ThermoFisher Scientific, Waltham, USA), followed by quantification using the JustTLC software (Sweday, Lund, Sweden).

Proteins were further purified by size-exclusion chromatography (SEC) using an ÄKTA pure 25 system (GE Healthcare Lifesciences) on a HiLoad 16/600 Superdex 75 pg column equilibrated in 25 mM Tris-HCl, 100 mM NaCl, pH 8.3. Protein concentrations were determined by measuring absorbance at 280 nm using theoretical molar extinction coefficients calculated with the ExPASy ProtParam tool.

### PETase candidate initial evaluation

Initial PET hydrolysis screening was performed using amorphous polyethylene terephthalate (aPET) film (Goodfellow Cambridge Ltd.; product GF25214475). Films were punched into coupons of approximately 8.5 mg and incubated in 1.5 mL tubes containing 1 mL of 100 mM sodium phosphate buffer (pH 8.0) supplemented with 500 nM (14.9 μg mL^-1^) PETcand1 (PET-KR1) or PETcand15 protein. Reactions were incubated at 30 °C under constant stirring at 300 rpm for 3 d. Following incubation, tubes were allowed to settle, and the supernatant was analyzed directly. Total aromatic soluble products (TPA, MHET, BHET, and higher soluble oligomers) were quantified by measuring absorbance at 242 nm using a Nanodrop 1000 spectrophotometer (Thermo Scientific). Background signal from time-zero controls was subtracted, and product concentrations were reported as TPA equivalents (TPAeq) calculated from a TPA standard curve. Thermal stability was assessed by differential scanning fluorimetry (DSF) through determination of the melting temperature (*T_m_*) of each candidate PETase. A 5000× SYPRO Orange stock was diluted in dH2O to 50×, and 2.5 μL of this working dye was mixed with 22.5 μL of Tris–HCl (pH 8.3) containing 10 μg PET_candX_ enzyme (or buffer blank) in a white, clear PCR plate (final reaction volume 25 μL; final dye 5×). Fluorescence was monitored on a CFX96 Touch Real-Time PCR System (Bio-Rad) in the FRET channel during a temperature ramp from 25 °C to 100 °C at 1 °C min^−1^. The *T_m_* was defined as the temperature at the minimum of the negative first derivative of the melting curve, as calculated in Bio-Rad CFX Maestro.

### Biochemical characterization of PET-KR1

Among the four enzyme candidates, PETcand1 was selected for further study and hereafter referred to as PET-KR1, whereas PETcand15 was renamed PET-KR2. Initial biochemical characterization of PET-KR1 was performed by quantifying esterase activity using the proxy substrate *p*-nitrophenyl butyrate (pNPB) via spectrophotometric detection of released *p*-nitrophenol (pNP) at 410 nm.

To determine the temperature optimum of PET-KR1 esterolytic activity, reactions (200 μL total volume) were prepared in 100 mM sodium phosphate buffer (pH 8.0) preincubated at 30–90 °C. SEC-purified PET-KR1 (5 μL) was added to 190 μL of pre-equilibrated buffer, and reactions were initiated by adding 5 μL pNPB (40 mM stock in DMSO), yielding a final enzyme concentration of 1.93 nM (57.5 ng mL^-1^) and 2.5% (v/v) DMSO. After 1 min incubation at the test temperature, absorbance at 410 nm was recorded using a Spectrostar Nano microplate reader (BMG Labtech) with the reader chamber set to 30 °C. The 1-min time point was chosen based on preliminary time-course measurements to fall within the linear range of pNP formation. Non-enzymatic hydrolysis (blank) was subtracted, and activity was reported as relative activity (%), normalized to the maximum value observed.

For determination of the pH optimum of PET-KR1, esterolytic activity was assayed over pH 5–9 using the pNPB esterase assay. Buffer systems (all 100 mM) were citrate–phosphate (pH 5–6), sodium phosphate (pH 6–8), and Tris–HCl (pH 8–9). Reactions (200 μL total volume) were prepared by adding 5 μL SEC-purified PET-KR1 to 190 μL of the respective buffer and initiating the reaction with 5 μL of pNPB (40 mM in DMSO) stock (final enzyme concentration 1.93 nM/57.5 ng mL^-1^; final DMSO 2.5% v/v). Absorbance at 410 nm was monitored at 30 °C for 10 min, and activities were calculated from the initial linear portion of the reaction curves after subtraction of the no-enzyme blanks. Activity was expressed as relative activity (%), normalized to the highest observed value.

For alkaline conditions, PET hydrolysis was assayed instead of the esterase assay to avoid autohydrolysis of the chromogenic substrate at high pH. Reactions were performed in sodium phosphate (pH 8.0), Tris–HCl (pH 8.0–9.0), or sodium carbonate (Na_2_CO_3_, pH 9.5) buffers. aPET film coupons (∼8.5 mg) were incubated in 1 mL of the respective buffer containing 456 nM (13.6 μg mL^-1^) SEC-purified PET-KR1 in 1.5 mL microcentrifuge tubes at 30 °C and 300 rpm for 24 h. Soluble aromatic products released during PET depolymerization were assessed by UV absorbance at 242 nm using a NanoDrop 1000 spectrophotometer (Thermo Scientific); signals were baseline-corrected by subtraction of the corresponding time-zero (t = 0) absorbance and expressed as relative activity (%), normalized to the highest blank-corrected value.

### PET depolymerization kinetics and product analysis

To evaluate the PET depolymerization profile of PET-KR1, reactions were performed at 40, 45, 50, 55 and 60 °C for up to 168 h, with sampling at 1, 2, 3, 4, 5, 6, 8, 10, 24, 48, 72, 96, and 168 h. Reactions (1 mL total volume) contained an aPET film coupon (8.5 mg) and were initiated by adding 26 μL SEC-purified PET-KR1 to 974 μL of 100 mM sodium phosphate buffer (pH 8.3) in 1.5 mL microcentrifuge tubes (final enzyme concentration 500 nM), followed by incubation at the indicated temperature with shaking at 300 rpm. At each time point, soluble aromatic products were quantified by measuring absorbance at 242 nm (NanoDrop), and signals were baseline-corrected by subtraction of the corresponding time-zero reading. Absorbance values were converted to TPA_eq_ using a TPA standard curve, and specific activity (μmol_products_ h^-1^ mg ^-1^) was calculated from the initial rates of the linear region of product formation when linearity was observed; for conditions that plateaued early, specific activities were not determined. Final specific productivity (μmol_products_ h^-1^ mg ^-1^) was calculated at 168 h from the endpoint TPA_eq_ values and compared across temperatures.

PET film depolymerization products were quantified spectrophotometrically as TPAeq, providing a rapid method for comparative kinetic screening across multiple temperatures. UV-based quantification of soluble aromatic PET hydrolysis products has previously been shown to correlate well with HPLC-derived measurements for relative activity comparisons and kinetic profiling ^56^. For the following cryo-milled PET experiments, product concentrations were determined by HPLC to enable individual quantification of TPA, MHET, and BHET and analysis of product distributions.

To evaluate the effects of temperature and PET morphology on the depolymerization activity of PET-KR1, reactions were conducted at 45, 50, 60, and 65 °C for up to 168 h. The same buffer was used with reactions (1 mL total volume) containing 8.5 mg of cryo-milled amorphous PET (aPET) powder (particle size < 500 μm) and were initiated by the addition of purified PET-KR1 fraction corresponding to 500 nM (14.9 μg mL^-1^) at the final reaction volume. Separate reaction tubes were prepared for each sampling time point and incubated under the specified conditions. At each time point, a single tube was removed from incubation for analysis.

At each time point, a defined volume of supernatant from the reaction tube was carefully withdrawn, appropriately diluted and filtered (0.20 μm). Depolymerization products were quantified by High-Performance Liquid Chromatography (HPLC) using a Macherey-Nagel NUCLEOSIL 100-5 C18 HD column (Dueren, Germany) at 40 °C and an isocratic mobile phase at a flow of 1 mL/min consisting of solvent A (Milli-Q water with 0.1% (v/v) trifluoroacetic acid; VWR, Radnor, PA) and solvent B (acetonitrile; Merck, Darmstadt, Germany) at an 80:20 (v/v) ratio. Standard curves were constructed using TPA, MHET and BHET (Sigma, Saint-Louis, USA) at known concentrations.

The total product release was calculated according to the following equation (Eq.1):

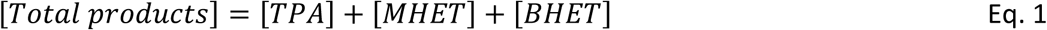

To compare product distributions among conditions, the following metrics were calculated based on the molar concentration of products, according to Equations 2 and 3:

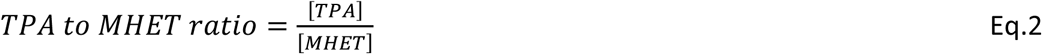

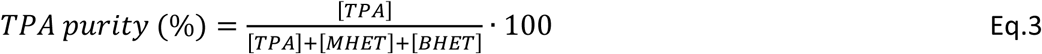

### Docking of PET tetramer

For docking analysis, a PET-tetramer model substrate (2-HE(MHET)_4_) was constructed from the SMILES string “OCCOC(=O)c1ccc(cc1)C(=O)OCCOC(=O)c1ccc(cc1)C(=O)OCCOC(=O)c1ccc(cc1)C(=O)OCCOC(=O)c 1ccc(cc1)C(=O)OCCO” and energy-minimized using the steepest descent algorithm with the MMFF94 force field (100 steps) in Avogadro (v1.2.0) ^57^. The PET-KR1 structure predicted by ColabFold (see Bioinformatic Screening section) was used as the receptor model. Protein and ligand structures were prepared using AutoDock Tools (v.1.5.7) ^58^, where polar hydrogens were added and Kollman charges were assigned to the protein. The ligand was assigned Gasteiger charges and converted to pdbqt format. As the oligomers of PET have very high conformability, compared to the post-consumer PET in which the chains do not exhibit freedom of movement ^59^, torsional rotations along ester bonds (between the aromatic ring carbon and carbonyl carbon, and between the carbonyl carbon and ester oxygen) were constrained to prevent intramolecular collapse during docking. Docking simulations were performed using AutoDock Vina (v1.1.2) ^60^ with a cubic grid box encompassing the entire protein (grid center: x = −1.052, y = −0.658, z = −0.981; box size: 68 × 68 × 68 Å). The exhaustiveness parameter was set to 25 and the energy range to 4 kcal mol⁻¹. For each docking run, nine poses were generated, and calculations were performed in triplicate. The top docking pose with the lowest predicted binding energy and a catalytically relevant orientation relative to the active site was selected for subsequent molecular dynamics simulations.

### Molecular dynamics simulations

Two types of systems, PET-KR1 (apo) and PET-KR1 in complex with PET-tetramer (holo) were employed for the molecular dynamics (MD) simulations using the GROMACS (v.2024.4) software ^61^. The systems were generated using a TIP3P water system and the CHARMM36 force field ^62^. The ligand parameters were prepared based on the CHARMM36 force field using the CGenFF server ^63^ and the protein-ligand complex was generated by combining the corresponding topologies and atom coordinates. A cubic box was centered around the protein and protein-ligand complex respectively with at least 1.2 nm distance around the molecules and the systems were solvated with water. Following solvation, the total system charges were neutralized by the addition of sodium and chloride ions. Subsequently, the systems were energy minimized using the steepest descent algorithm with 5X10^6^ steps and a 0.01 step size until a maximum force lower than 100 kJ/mol was achieved. Following energy minimization, the systems underwent stepwise equilibration. First, NVT equilibration was performed for 100 ps at 300 K using the V-rescale thermostat (τ_t = 0.1 ps; 2 fs timestep). This was followed by NPT equilibration for 100 ps at 300 K and 1 bar using the V-rescale thermostat (τ_t = 0.1 ps) and C-rescale barostat (τ_p = 2.0 ps; 2 fs timestep). During both equilibration stages, harmonic position restraints (1000 kJ mol⁻¹ nm⁻²) were applied to the PET tetramer ligand heavy atoms (defined via indexing a restrained ligand topology) to maintain proximity to the initial docked orientation. Production MD simulations were then performed for 500 ns per system in the NPT ensemble at 300 K and 1 bar, using the V-rescale thermostat (τ_t = 0.1 ps) and Parrinello–Rahman barostat (τ_p = 2.0 ps). Bonds involving hydrogen atoms were constrained using the LINCS algorithm, enabling a 2 fs integration timestep. A Verlet cutoff scheme applied 1.2 nm cutoffs for short-range electrostatics (PME; pme_order = 4; fourierspacing = 0.16 nm) and van der Waals interactions (force-switch 1.0–1.2 nm). All ligand restraints were removed for production runs. Each system (apo and holo) was simulated in triplicate, with independent replicates initiated from distinct atomic velocity distributions during system preparation.

Periodic boundary conditions were applied in all directions, and center-of-mass motion was removed at each timestep. System stability was evaluated by root-mean-square deviation (RMSD) of backbone atoms using least-squares fitting, as implemented in the gmx rms module. Residue-level flexibility was assessed via root-mean-square fluctuation (RMSF) analysis using the gmx rmsf module. Binding free energies (ΔG_bind) were calculated for the holo protein complex using gmx_MMPBSA (v1.5.1) ^64,65^ for each triplicate production trajectory and total binding energies were averaged across replicas. Protein–ligand interactions were further analyzed by calculating distances between protein residues and the PET tetramer across holo simulations using MDAnalysis ^66,67^. Residues were considered in contact with the ligand if any atom distance was ≤ 4.5 Å. Interaction occupancy was defined as:

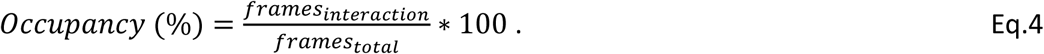

Finally, the residues with occupancy (%) ≥ 30% across MD replicas were retained and the Average occupancy (%) and Average distance from the ligand were calculated to highlight putative PET-binding regions.

### Rational disulfide engineering of PET-KR1

To further explore whether the thermostability of PET-KR1 could be improved for enhanced PET depolymerization at elevated temperatures, a rational disulfide-bond engineering strategy was applied. Two independent disulfide-prediction platforms, Disulfide by Design 2 and the Maestro web server, were used to identify candidate residue pairs in the PET-KR1 AlphaFold-predicted model ^68,69^. Four putative disulfide pairs were prioritized for experimental testing based on literature-supported positions in related PET hydrolases, consensus between the two prediction tools, and the structural feasibility of the proposed cysteine substitutions: R1, N206C/S260C; R2, E15C/P220C; R3, G6C/P20C; and R4, A142C/V168C. These candidates were selected as a small pool of putative stabilizing disulfide bonds for downstream validation.

The R1 variant was first generated as a single mutant and used as the reference engineered construct for comparison with the wild-type enzyme. In parallel, combinatorial variants were designed to test whether additional predicted disulfides could act synergistically with R1, including R1+R2, R1+R3, and R1+R4. The corresponding cysteine-substitution constructs were gene-synthesized and cloned into the pET-29b(+) vector (Twist Bioscience), similarly to the wild-type construct, and recombinant proteins were expressed under the same conditions.

### Data Analysis

All experimental data are reported as mean ± standard deviation (SD). A minimum of three independent experiments were performed for all assays except stated otherwise. Unless otherwise stated, data are reported as mean ± standard deviation (SD). For the initial aPET film degradation activity and *Tm* evaluation, a minimum of three independent experiments, each comprising technical triplicates were performed. pNPB esterase pH optimum assays were performed in nine independent experiments, each comprising technical triplicates. pNPB esterase temperature optimum and aPET film pH optimum assays were performed in three independent experiments, each comprising technical duplicates. The aPET film degradation assays were performed in technical triplicate measurements while the HPLC-based aPET powder depolymerization experiments were performed as technical duplicates. Specific activity was determined in the initial linear region of product formation, and specific productivity were calculated from endpoint (168 h) TPA-equivalent (film kinetics) or total HPLC-measured (powder kinetics) products values. MM/PBSA binding free energies are reported as the mean ± SD across three independent MD replicas. Comparisons between conditions and between enzymes are presented descriptively; no formal hypothesis tests were applied, given the number of independent replicates. Data were processed and plotted in GraphPad Prism v9.0.0.

## Results and discussion

### Targeted screening of 277 plastic-associated metagenomes yields 21 high-confidence PETase candidates

Analysis of 277 metagenomes from environments exposed to plastics using *ProteoSeeker* ^37^ identified over 77 million predicted proteins (Figure 1A). At the time of the analysis (November 2024), no dedicated PETase Pfam domain existed, necessitating use of custom Hidden Markov Models (HMMs) and proxy domains—cutinase (PF01083), chlorophyllase (PF07224, PF12740), and dienelactone hydrolase (PF01738)—consistent with contemporary strategies reported in very recently published studies ^30,32,33,70^. Consequently, seek mode (Pfam-targeted) and taxonomy mode (domain-agnostic) outputs from *ProteoSeeker* were pooled without domain exclusion, yielding 54 candidates bearing canonical PETase Type I/II catalytic motifs (Gly-His-Ser-Met-Gly-Gly-Gly-Gly or Gly-Trp-Ser-Met-Gly-Gly-Gly-Gly). All 54 candidates passed length filtering (250–350 amino acid residues). C-terminal disulfide analysis—requiring ≥2 cysteines at specific positions validated by MAFFT MSA with characterized PETases—identified 30 candidates. CD-HIT (100% identity) dereplication yielded 21 unique sequences. TM-Align superposition of AlphaFold models vs LCC (PDB:4EB0) confirmed all 21 exhibited TM-scores ≥0.9 (mean 0.95±0.01), demonstrating highly conserved α/β-hydrolase folds regardless of Pfam status.

**Figure 1.**
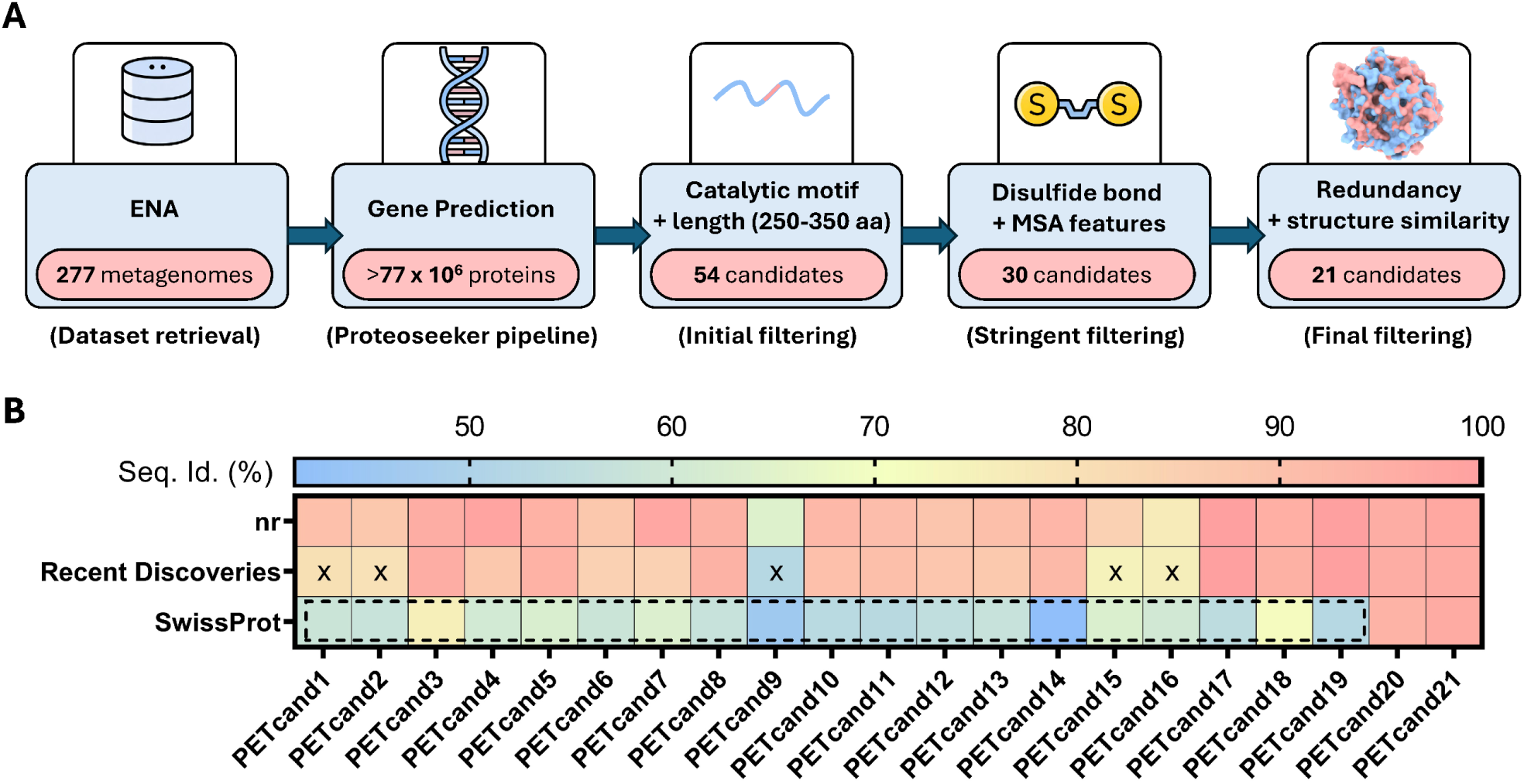
Discovery pipeline & Sequence identity heatmap of putative PETases from plastic metagenomes. **(A)** Schematic overview of the *ProteoSeeker*-based discovery workflow: 277 plastic-associated metagenomes were retrieved from ENA, translated via gene prediction (>77 million predicted proteins), and filtered by PETase catalytic motifs and length (250–350 aa) to yield 54 candidates; subsequent stringent filtering based on disulfide-bond/MSA features reduced this set to 30 candidates, and final redundancy removal plus structural similarity filtering yielded 21 non-redundant candidates. **(B)** Maximum % identity values for PETcand1–21 via BLASTP against Swiss-Prot (2025_12), nr, and PETase sets from Seo et al. (2025), Norton-Baker et al. (2025), and Wu et al. (2025); all hits ≥82% coverage. Color scale: light blue (∼41%) to light red (100%). Dashed lines denote low Swiss-Prot identity candidates; × marks five novel sequences (52–80% identity).

This combined validation filtering ensured candidate robustness independent of evolving domain annotations. Satisfyingly, re-analysis of these 21 final candidates using current HMMER databases revealed universal PETase domain assignment. PF12740—originally annotated as “Chlorophyllase2” during our initial November 2024 screening—now consistently identifies as “PETase/PET hydrolase-like” across all sequences: the original 12 seek-mode hits plus the 9 taxonomy-mode hits lacking user-specified domains. This demonstrates recent Pfam model evolution incorporating PETase-specific sequence features discovered post-analysis.

However, the presence of a PF12740 domain is not sufficient on its own. Although this annotation indicates similarity to the broader PETase-like family, proteins annotated with this domain very often lack one or more essential determinants of *bona fide* activity, including the complete catalytic motif and the C-terminal disulfide bond. Our multi-step filtering therefore, was essential to distinguish biochemically relevant PETases from domain-positive false positives.

### Plastic-associated metagenomes yield candidates spanning established PETase subtypes and hybrid architectures

BLASTP against Swiss-Prot (2025_12) revealed 19 candidates (PETcand1–19) with modest identity (41–76%) to characterized PETases, while PETcand20–21 exceeded 93% (Figure 1). nr hits were predictably higher for most, except PETcand9 and PETcand16 (max 63.8% and 76.6%), reflecting nr’s abundance of predicted homologs versus the curated, biochemically validated entries of Swiss-Prot. The primary sequences of PETcand1–21 are reported in Supplementary Figure S1.

Since recent 2025 PETase discoveries (e.g., Seo *et al*., Norton-Baker *et al*. and Wu *et al*. ^24,30,31^) are incompletely annotated in Swiss-Prot, an additional 2,575 sequences from these studies were screened. Notably, only five candidates (PETcand1,3,9,15,16; ≤80% max %ID, marked by ×) remain novel against this expanded set, underscoring the rapid expansion of PETase sequence space and need for experimental validation (Figure 1B).

Maximum-likelihood phylogenetic analysis of the selected 21 putative PETases and 13 biochemically characterized PETases was performed. α/β-hydrolase references resolved distinct PETase-linked radiation within the superfamily, rooted against classical outgroups including dienelactone hydrolases (DLH; Figure 2). While PETases do not form a strictly monophyletic clade excluding all other hydrolases, the topology reveals well-supported subclusters corresponding to established subtypes.

**Figure 2.**
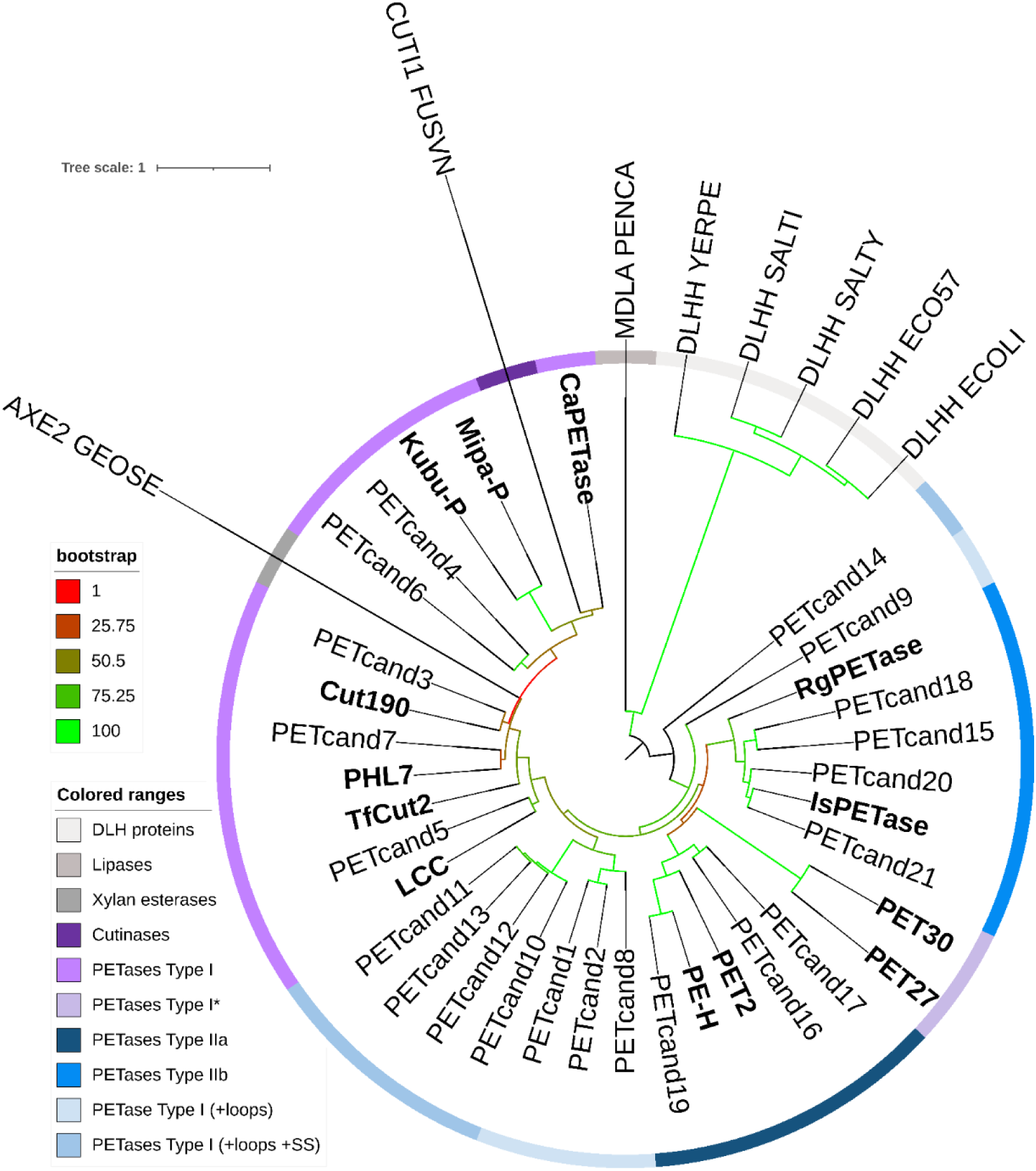
Phylogenetic analysis of PETases and related α/β-hydrolases. Maximum-likelihood tree of the 21 putative PETases from plastic-associated metagenomes identified in this work, 13 biochemically characterized PETases, and representative α/β-hydrolases. PETases are colored by established functional subtypes: Type I (purple), Type I* (light purple), Type IIa (dark blue) and Type IIb (blue). Metagenome-derived candidates are shown in black (this work) while characterized PETases are shown in bold. Cutinase (CUT1) and acetyl xylan esterase (AXE2) cluster within the broader PETase-associated radiation, whereas dienelactone hydrolases (DLH) and lipases were included as distant outgroups. Bootstrap values (1,000 replicates) are shown at key nodes.

Within the PETase-associated radiation, several well-supported subclusters are observed: Type I (purple; GHSMGGGG motif, single C-terminal disulfide) containing the well-known enzymes LCC, Mipa-P etc., plus GWSMGGGG enzymes (Kubu-P, CaPETase) that cluster with canonical Type I topology. A second radiation features extended loop regions neighboring Type I, encompassing Type IIa (dark blue), Type IIb (blue), and Type I* (light purple) subgroups—all bearing the GWSMGGGG motif.

The 21 candidates identified here distribute across this PETase radiation rather than a single subtype. Notably, PETcand1–2 and PETcand9–14 occupy intermediate positions within the PETase-associated radiation, combining Type I GHSMGGGG motifs with Type II-like extended loops and secondary disulfides—combinations of sequence and structural features typically associated with distinct PETase subtypes (Figure 2).

Consistent with prior reports that some cutinase- and esterase-like hydrolases exhibit low-level or promiscuous activity toward synthetic polyesters, reference sequences (CUT1, AXE2) embed within the PETase-linked cluster rather than distant outgroups ^71,72^. Critically, no metagenomic candidates associate with classical outgroups, confirming their PETase relatedness. This distribution—spanning established subtypes plus putative hybrids—demonstrates plastic-associated metagenomes as rich sources of putative PETase diversity.

To identify key sequence features of PETases, all putative enzymes were aligned with characterized reference PETases using MAFFT and visualized using the ESPript server (Figure S1), while a focused alignment of selected candidates appears in Figure 3. Generally, PETases featuring a single C-terminal disulfide bond and the characteristic GHSMGGGG motif are classified as Type I. Conversely, enzymes bearing the GWSMGGGG motif with extended α2-α3 and β8-α6 loops represent either Type I* or Type II subtypes. Type I* retain a single (slightly repositioned) C-terminal disulfide, whereas Type II possess an additional disulfide connecting β7 beta sheet with η4 alpha helix. Type IIb enzymes exhibit Ser at the position homologous to Ser238 in IsPETase, while Type IIa harbor Tyr, Phe, or Trp at this site.

**Figure 3.**
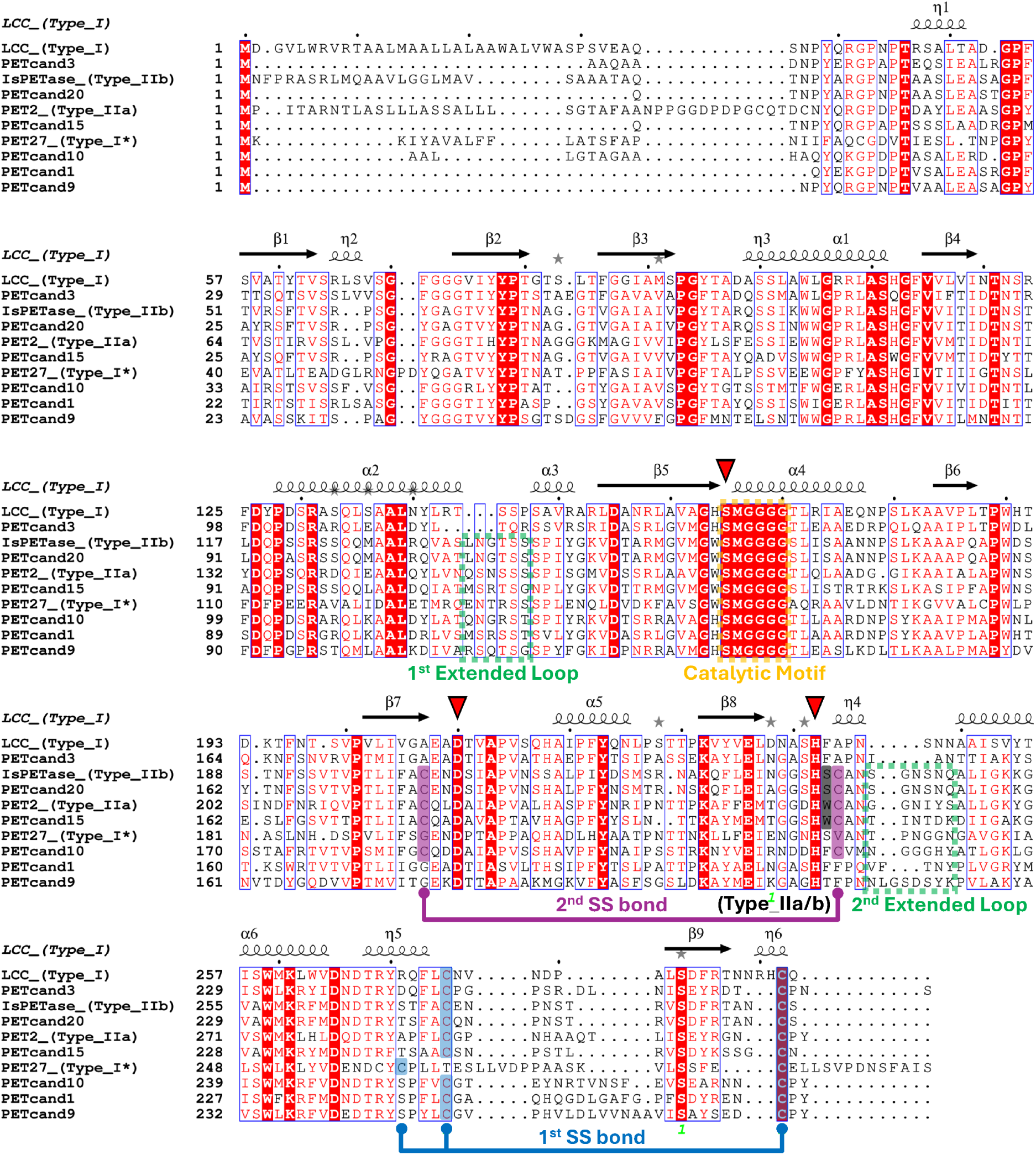
Multiple sequence alignment of metagenomic PETase candidates (this work) with characterized Type I, IIa, IIb, and I* reference PETases. Secondary structure elements derived from LCC crystal structure (PDB: 6THS). Key diagnostic features annotated: Ser-His-Asp catalytic triad (red triangles), C-terminal disulfides (purple/blue lines), α2-α3 and β8-α6 loops characteristic of Type II/I* enzymes (green boxes), catalytic motifs (GHSMGGGG/GWSMGGGG; orange), and Type IIb-defining Ser238 (black). Protein sequences aligned using MAFFT and visualized with ESPript server.

Intriguingly, the unclassified enzymes Kubu-P and CaPETase—which possess the GWSMGGGG motif but lack other Type II characteristics—cluster with Type I PETases in our phylogenetic analysis (Figure 2). PETcand3–7 align as canonical Type I sequences within the corresponding clade. PETcand15–21 appropriately position near Type II references, with Type IIb candidates generally accurate except PETcand15 (Trp) and PETcand18 (Phe) at the subtype-defining position; PETcand16,17,19 cluster near Type IIa. Notably, PETcand1,2,8,9 bear the Type I GHSMGGGG motif alongside Type II-like extended loops, while PETcand10–14 additionally possess the second disulfide. These hybrid sequences predominantly cluster between Type I and Type II/I* clades, consistent with their mixed features—except PETcand9 and PETcand14 near the root. Similar feature combinations have been reported in previously characterized PET hydrolases, underscoring the structural variability within the family. PETcand10–14 exhibit high similarity to low-expression, negligible PET hydrolysis clusters, whereas PETcand8 homologs show expression, and PETcand1,2,9 represent relatively novel sequences.

The combined analyses prioritized PETcand1 and 9 (low-identity Type I-like hybrids with extended loops) and PETcand15 and 16 (canonical Type II with novel sequence profiles) for experimental validation (Figures 1–3). PETcand2 was excluded due to ≥90% sequence identity with PETcand1. These four candidates—spanning hybrid and established subtypes from plastic-associated metagenomes—were selected for recombinant expression, PET-degrading activity verification, and biochemical characterization.

### Functional screening of metagenomic sequences reveals two novel PETases

To experimentally validate PET depolymerization activity, four prioritized metagenome-derived PETase candidates (PETcand1, PETcand9, PETcand15, and PETcand16) were selected for recombinant production based on their sequence novelty. Genes encoding the mature enzymes (signal peptide removed) were codon-optimized for *E. coli*, cloned in pET-29b (+) vectors to introduce a C-terminal His tag, and expressed in *E. coli* SHuffle T7 Express to support cytosolic disulfide bond formation. Recombinant proteins were purified by immobilized metal affinity chromatography (IMAC) followed by size-exclusion chromatography (SEC).

Of the four candidates, only PETcand1 and PETcand15 yielded sufficient soluble protein and were obtained as SEC-purified preparations for downstream screening. PETcand9 was detected only in the insoluble fraction, whereas PETcand16 was undetectable in both soluble and insoluble fractions. Initial PET hydrolysis activity was assessed using amorphous PET (aPET) film coupons (∼8.5 mg) incubated with 500 nM (14.9 μg mL^-1^) enzyme in 100 mM sodium phosphate buffer (pH 8.0) at 30 °C and 300 rpm for 3 d. Released soluble aromatic products (TPA, MHET, BHET, and soluble oligomers) were quantified by absorbance at 242 nm, baseline-corrected to time-zero controls, and reported as terephthalic acid equivalents (TPAeq) based on a TPA standard curve.

Under these screening conditions, both enzymes produced detectable soluble products, with PETcand1 showing 3.1-fold higher product release than PETcand15 (Figure 4A). In parallel, differential scanning fluorimetry (DSF) indicated higher apparent thermostability for PETcand1 (*T_m_* = 66.5 °C) than for PETcand15 (*T_m_* = 53.6 °C) (Figure 4A and Figure S2). PET depolymerization is promoted by a combination of high catalytic activity and sufficient thermostability to sustain turnover over extended reaction times, particularly when operating close to the glass-transition temperature of PET (Tg ≈ 70 °C), where increased polymer-chain mobility can improve enzyme access to ester bonds. Based on its favorable balance of catalytic output and thermostability in our initial screening, PETcand1 was selected for detailed biochemical and process-level characterization and is hereafter referred to as PET-KR1.

**Figure 4.**
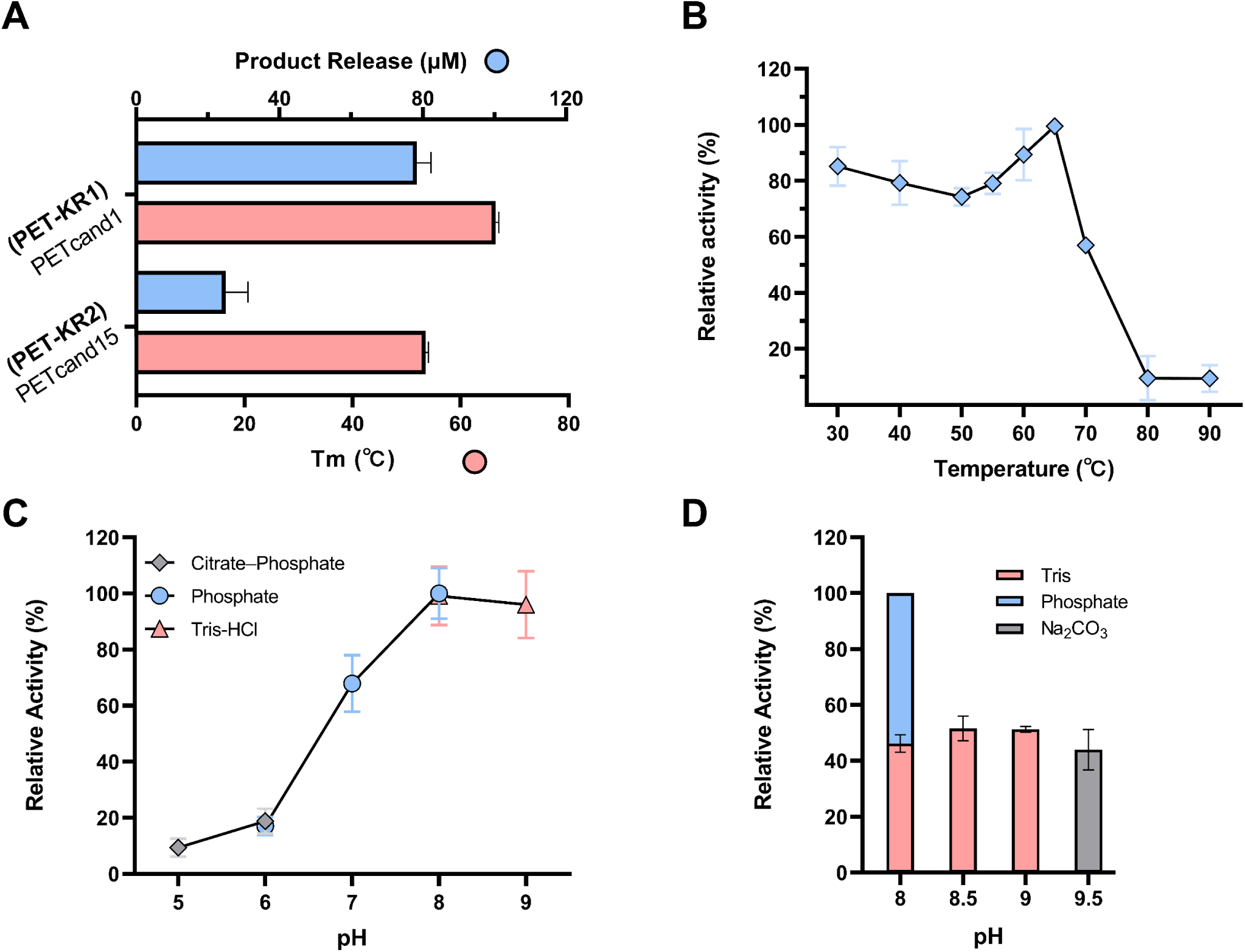
Initial screening and biochemical characterization of PET-KR1. **(A)** Initial PET hydrolysis screening of SEC-purified PETcand1 (PET-KR1) and PETcand15 using amorphous PET (aPET) film coupons at 30 °C for 3 days; soluble aromatic products released to the supernatant were quantified by UV absorbance at 242 nm and reported as TPA equivalents (TPAeq) (blue bars), and apparent melting temperatures (*T_m_*) were determined by differential scanning fluorimetry (DSF) using SYPRO Orange (pink bars). **(B)** Temperature dependence of PET-KR1 esterase activity (30–90 °C, pH 8.0) measured using *p*-nitrophenyl butyrate (pNPB) and expressed as relative activity normalized to the maximum. **(C)** pH dependence of PET-KR1 esterase activity (pH 5.0–9.0) measured with pNPB across the indicated buffer systems (100 mM citrate–phosphate, sodium phosphate, or Tris–HCl) and expressed as relative activity normalized to the maximum. **(D)** pH dependence of PET-KR1 PET hydrolysis under alkaline conditions (pH 8.0–9.5) assessed by aPET depolymerization and quantification of soluble aromatic products at 242 nm in the indicated buffers (100 mM sodium phosphate, Tris–HCl, or Na_2_CO_3_) activities are shown relative to the maximum.

### Temperature and pH dependence of PET-KR1 activity

To understand better the functional window of PET-KR1, its esterase activity was quantified using the proxy substrate pNPB across a broad temperature range (30–90 °C) at pH 8.0. PET-KR1 maintained high relative activity between 30 and 65 °C, reaching a maximum at 65 °C. Then, it exhibited a steep activity decline at ≥70 °C (Figure 4B), consistent with thermal inactivation as assay temperatures approach the protein’s unfolding transition. This trend is in agreement with the aforementioned DSF analysis, which had revealed a *T_m_* of 66.5 °C (Figure 4A), where *T_m_* corresponds to the midpoint of the thermal unfolding transition (i.e., ∼50% of the protein population unfolded under the assay conditions) ^73^. As the folded fraction decreases and unfolded species accumulate, catalytic activity typically drops due to loss of the active-site geometry and, in many cases, irreversible denaturation pathways such as aggregation ^74,75^. Together, the close agreement between the temperature optimum (65 °C), the DSF-derived *T_m_* (66.5 °C), and the sharp activity loss above 70 °C supports thermostability as a key factor limiting PET-KR1 performance at elevated temperatures.

The pH dependence of PET-KR1 esterase activity was assessed using the soluble proxy substrate pNPB from pH 5.0 to 9.0 in three 100 mM buffer systems (citrate–phosphate, sodium phosphate, and Tris–HCl). PET-KR1 activity increased with pH and plateaued at pH 8–9 (Figure 4C), indicating a preference for neutral-to-alkaline conditions in the pNPB assay. Because alkaline conditions can increase non-enzymatic hydrolysis of chromogenic esters, PET depolymerization (aPET film) was used to directly evaluate performance at higher pH values (pH 8.0–9.5) in sodium phosphate, Tris–HCl, and Na_2_CO_3_ buffers (Figure 4D). In this substrate-relevant assay, maximal activity was observed at pH 8.0 in sodium phosphate, whereas the same nominal pH (8.0) in Tris–HCl yielded lower activity (∼46% relative), pointing to buffer-identity effects rather than pH alone. This interpretation is supported by prior work showing that Tris can inhibit PET hydrolysis for cutinase-like PET hydrolases (e.g., LCC and TfCut2), consistent with interference/competition effects, which are more pronounced for polymeric substrates than for small soluble esters such as pNPB ^76^. Despite this buffer dependence at pH 8.0, PET-KR1 retained >40% relative PET hydrolysis activity across pH 8.5–9.5 in Tris and carbonate buffers, supporting robustness under mildly alkaline conditions. Thus, in line with Standardization guidelines for bacterial PET hydrolases, where lab-scale PET depolymerization assays are frequently pH-controlled near 7–8 using low-concentration phosphate or Tris buffers, 100 mM sodium phosphate buffer (pH 8.0) was selected for downstream experiments unless stated otherwise ^77,78^.

### PET film depolymerization kinetics for PET-KR1

To quantify PET-KR1 performance under process-relevant, prolonged reaction conditions, we monitored aPET film depolymerization at 40–60 °C for up to 168 h and quantified soluble products as TPA equivalents (TPAeq) over time (Fig. 5A). Importantly, this time-course experiment was designed as a process-oriented screen to identify operating conditions that balance early productivity and sustained activity on relevant PET substrate, rather than to define the enzyme’s intrinsic catalytic optimum on a soluble model ester. Total product accumulation profiles revealed strong temperature dependence, with substantial depolymerization observed at 40–50 °C, while reactions at 55–60 °C produced only minimal soluble products over the full-time course (Figure 5A, C). Specific activity, calculated from the first 4 h of product formation, peaked at 50 °C (∼2.7 μmol_products_ h^-1^ mg_enzyme_^-1^) (Fig. 5B), despite the higher temperature optimum (65 °C) measured in the short pNPB esterase assay (see Temperature and pH dependence of PET-KR1 activity section). This shift is consistent with thermostability limiting performance in the PET film assay: unlike the pNPB readout (1-min incubation), PET depolymerization requires sustained catalysis over hours to days on an insoluble, hydrophobic polymer surface and is therefore more sensitive to time-dependent thermal inactivation, particularly at elevated temperatures. Accordingly, the 50 °C reaction exhibited rapid early product release followed by a pronounced slowdown after 48 h, reaching a plateau and specific productivity of 0.57 μmol_products_ h^-1^ mg_enzyme_^-1^ by 168 h (Figure 5A, C). At 45 °C, product formation remained robust for longer and only began to decelerate after 96 h, ultimately reaching the highest specific productivity over the 168-h reaction (∼0.73 μmol_products_ h^-1^ mg_enzyme_^-1^) (Figure 5A, C). In contrast, at 40 °C, product accumulation proceeded more steadily across the entire time window without an obvious plateau, but with a lower final specific productivity (∼0.4 μmol_products_ h^-1^ mg_enzyme_^-1^) (Figure 5A, C). Together, these data indicate that PET-KR1 has high intrinsic esterolytic potential at elevated temperature (as suggested by the pNPB profile); yet, its effective PET depolymerization performance over multi-day reactions is constrained, suggesting that functional stability becomes limiting above ∼50 °C and, potentially, limited by surface accessibility of the PET film to the enzyme, highlighting thermostability as a key target for applying protein engineering to unlock higher-temperature PET hydrolysis.

**Figure 5.**
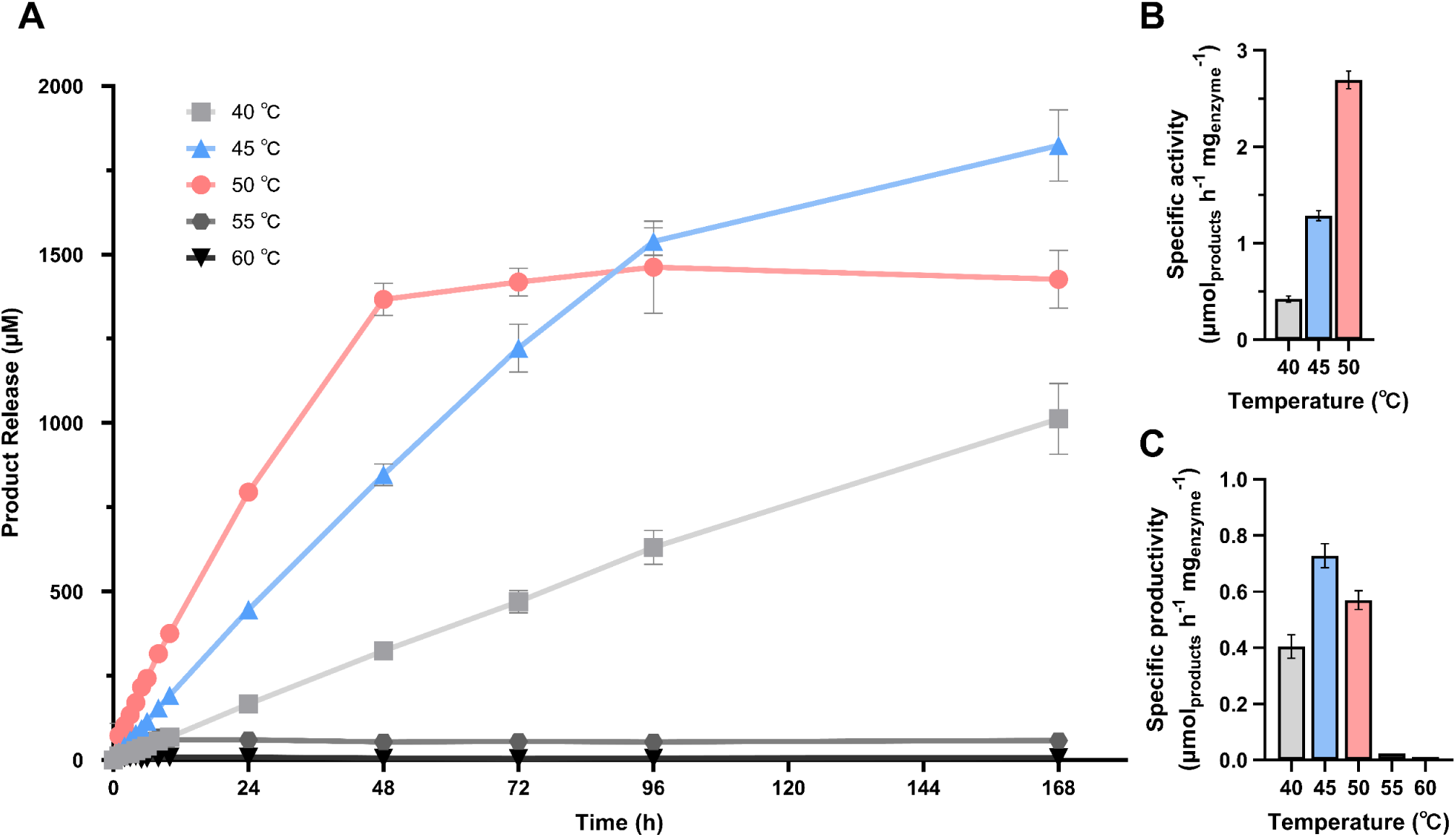
PET film depolymerization kinetics of PET-KR1 across temperature. **(A)** Time-course release of soluble aromatic products during depolymerization of amorphous PET (aPET) film by PET-KR1 at 40, 45, 50, 55, and 60 °C over 168 h. Soluble products were quantified by UV absorbance at 242 nm, baseline-corrected to the corresponding time-zero control and converted to TPAeq using a TPA standard curve. **(B)** Specific activity per mg of enzyme calculated from the linear region within the first 4 h of the reaction. **(C)** Final specific productivity of total products (end-point TPAeq) after 168 h as a function of temperature.

### PET powder depolymerization time courses reveal enhanced substrate accessibility and a shifted temperature apparent process optimum

To evaluate the effect of PET morphology on enzyme performance, PET-KR1 was incubated with cryo-milled amorphous PET (aPET) powder at 45, 50, 60 and 65 °C for up to 168 h, and soluble depolymerization products (TPA, MHET and BHET) were quantified by HPLC. In contrast to PET film depolymerization, where maximal specific productivity was observed at 45 °C and declined markedly at higher temperatures, aPET powder hydrolysis showed a clear apparent process optimum at 50 °C, reaching approximately 3.61 μmol_products_ h^-1^ mg_enzyme_^-1^ of total soluble products after 168 h (Figure 6A, C). By comparison, reactions at 45 °C accumulated 1.66 μmol_products_ h^-1^mg_enzyme_^-1^, whereas reactions at 60 °C reached an early plateau in product formation and final specific productivity of 1.8 μmol_products_ h^-1^ mg_enzyme_^-1^, while virtually no product formation was detected at 65 °C.

**Figure 6.**
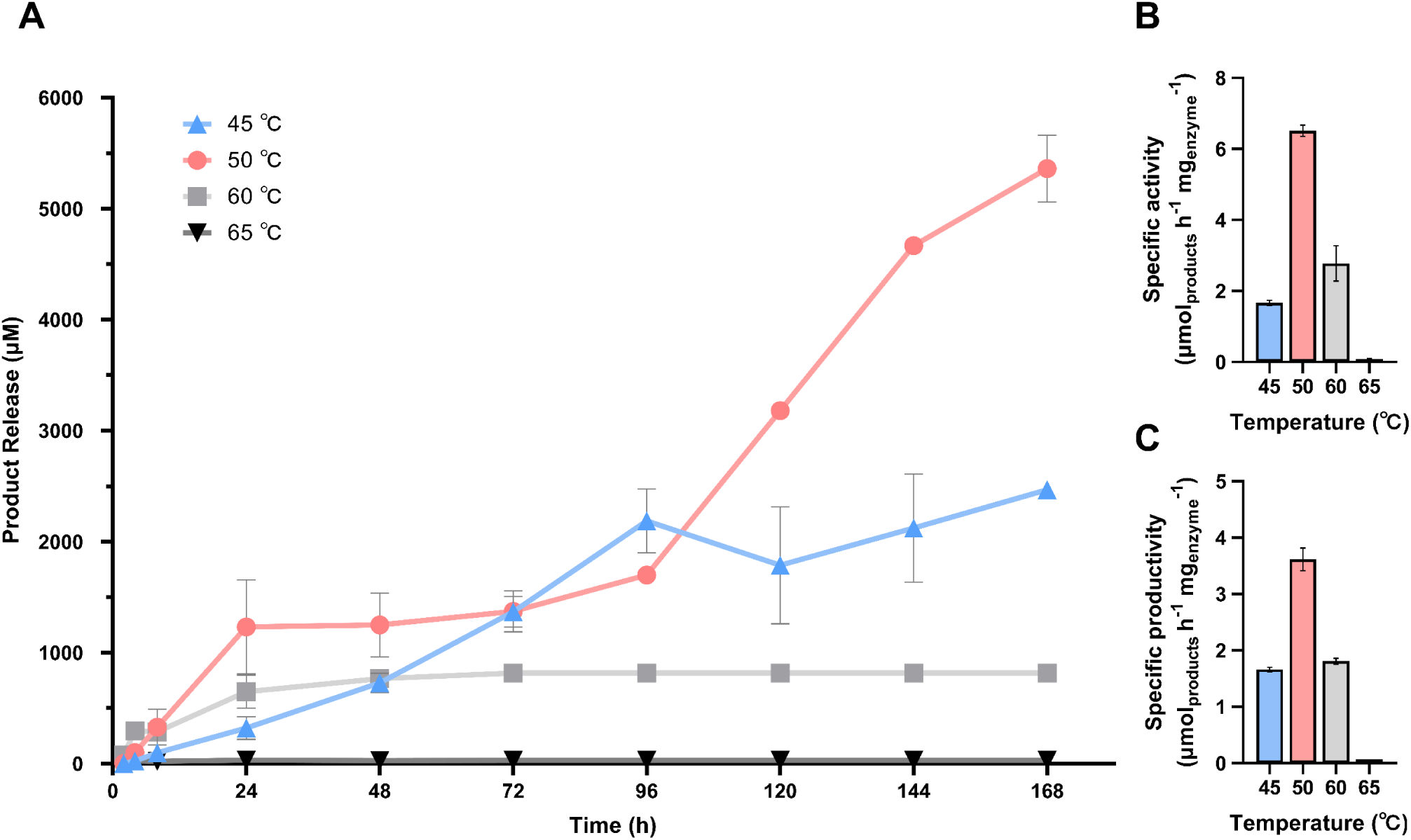
PET powder depolymerization product profile over different times and temperatures. **(A)** Time-course release of soluble aromatic products during depolymerization of amorphous PET (aPET) powder by PET-KR1 at 45, 50, 60 and 65 °C over 168 h. Soluble products were quantified by HPLC-UV and refer to the sum of produced TPA, MHET and BHET **(B)** Specific activity calculated from the linear region **(C)** Specific productivity after 168 h as a function of temperature.

The specific activities followed the same trend, increasing from 1.66 μmol_products_ h^-1^ mg_enzyme_^-1^ at 45 °C to nearly 6.51 μmol_products_ h^-1^ mg_enzyme_^-1^ at 50 °C before decreasing substantially at 60 °C (Figure 6B). The higher initial rates observed for PET powder relative to PET film likely reflect the substantially larger accessible surface area of the cryo-milled substrate, which facilitates enzyme adsorption and ester-bond cleavage. Notably, the initial specific rate at 50 °C exceeded that measured for aPET film under the same conditions (2.7 μmol_products_ h^-1^ mg_enzyme_^-1^), while the specific productivity was approximately 5-fold higher than the maximum obtained with aPET film (0.72 μmol_products_ h^-1^ mg_enzyme_^-1^ at 50 °C). These observations are consistent with substrate accessibility becoming a major limitation during film depolymerization.

The product accumulation profiles further revealed distinct kinetic behaviors. At 50 °C, product release accelerated after an initial lag phase and continued throughout the 168-h incubation, suggesting sustained catalytic activity and possible progressive access to newly exposed polymer surfaces. In contrast, the 45 °C reaction displayed slower but more stable product accumulation, while the 60 °C reaction rapidly approached a plateau, consistent with partial thermal inactivation of PET-KR1 despite the improved substrate accessibility of the powder. The absence of detectable depolymerization at 65 °C is consistent with the enzyme’s measured melting temperature (*T_m_* = 66.5 °C) and supports, once again, thermostability as the principal factor limiting performance at elevated temperatures. Although PET film and powder depolymerization were quantified using different analytical approaches, UV-based TPA-equivalent measurements have been shown to reproduce relative activity trends observed by HPLC and are widely used for kinetic screening of PET hydrolases ^56^. Therefore, the substantially higher productivity observed for cryo-milled PET is unlikely to be attributable solely to analytical methodology and therefore likely reflects a genuine effect of substrate morphology rather than an analytical artifact. Overall, the powder depolymerization experiments demonstrate that increasing substrate surface area substantially enhances PET-KR1 performance and shifts the apparent process optimum from 45°C for aPET film to 50 °C for aPET powder, highlighting the strong influence of substrate morphology on depolymerization efficiency.

To contextualise PET-KR1 within the PETase landscape, we performed a semi-quantitative comparison with the benchmark wild-type enzymes Mipa-P, Kubu-P, CaPETase, IsPETase, and LCC, based on data reported by Seo et al. (2025) ^24^ under closely related conditions (Table S1). Because substrate loading, PET morphology, agitation, enzyme concentration, and other unreported experimental details differ between studies, this comparison is intended for contextual benchmarking rather than strict kinetic equivalence. Under this framework, PET-KR1’s specific activity at 50 °C (6.5 μmol h⁻¹ mg⁻¹) is comparable to the highest specific activities reported for IsPETase and CaPETase, while its specific productivity at 50 °C (3.6 μmol h⁻¹ mg⁻¹) is the highest among the five reference wild-type enzymes. Its *Tm* of 66.5 °C is consistent with the lower-*Tm* benchmarks, though substantially below the highly engineered Kubu-P and LCC variants.

### Product distribution analysis reveals enhanced monomerization at elevated temperatures

To further investigate the PET-depolymerization behavior of PET-KR1, product distributions were analyzed using the TPA/MHET ratio and TPA purity metrics (Figure 7). Whereas total product release reflects overall depolymerization efficiency, these parameters provide insight into the extent of intermediate accumulation and monomer formation during PET hydrolysis.

**Figure 7.**
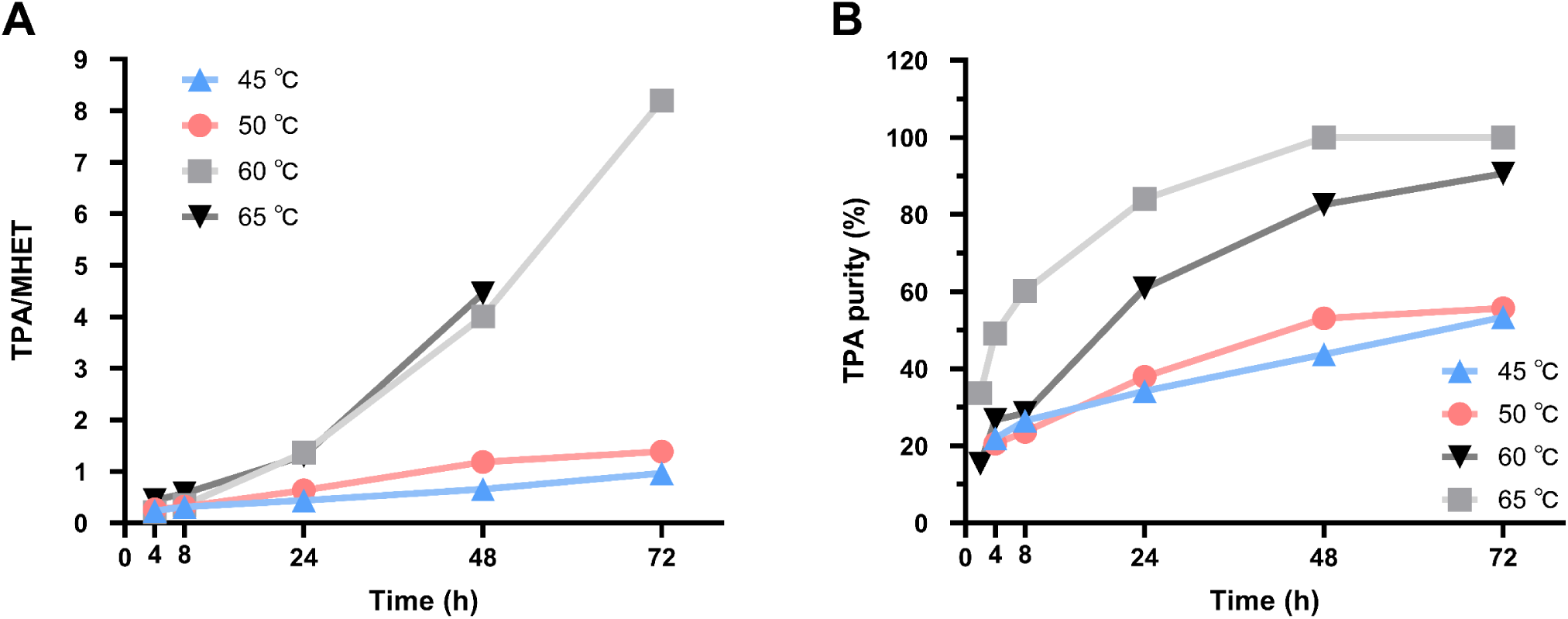
Product distribution analysis during PET hydrolysis. **(A)** TPA-to-MHET ratio and **(B)** TPA product purity from PET hydrolysis over time at different temperatures.

The TPA/MHET ratio of PET-KR1 increased progressively over time under all conditions tested, indicating gradual conversion of MHET into terephthalic acid (TPA) (Figure 7A). However, the magnitude of this increase was strongly temperature dependent. At 45 and 50 °C, the ratio remained below 1.5 after 72 h, indicating substantial accumulation of MHET relative to TPA. In contrast, reactions at 60 °C showed much higher TPA/MHET ratios, reaching approximately 8.2 after 72 h, while at 65 °C, MHET was fully depleted. Consistent with these observations, TPA purity increased continuously throughout the reaction and approached 90–100% at 60–65 °C, whereas only ∼55% TPA purity was observed at 45–50 °C after 72 h (Figure 7B). These results indicate that although overall product formation by PET-KR1 was lower at elevated temperatures (Figure 6), the soluble products released under these conditions were enriched in the final monomer TPA, indicating a shift toward more complete hydrolysis of released intermediates.

Interestingly, BHET, an upstream intermediate in the PET hydrolysis pathway, was not detected during PET depolymerization, suggesting that any BHET released by PET-KR1 may be rapidly converted to MHET before measurable accumulation occurs. This observation is consistent with a depolymerization profile that favors downstream monomer formation. Overall, these results suggest that PET-KR1 exhibits moderate depolymerization activity together with a favorable monomerization profile near its upper operational limit.

### Docking and molecular dynamics indicate stable PET binding in PET-KR1

To gain molecular insight into PET binding, a PET tetramer model substrate was docked onto the AlphaFold-predicted structure of PET-KR1. The top-ranked docking pose yielded a binding score of −9.2 kcal/mol and positioned the polymer within a shallow surface cleft, near the catalytic triad. The ester bond of the substrate was oriented in a catalytically relevant configuration, consistent with binding modes reported for previously characterized PETases. Molecular dynamics (MD) simulations were subsequently performed to evaluate the stability of the PET–PET-KR1 complex. Analysis of backbone RMSD and residue-level RMSF profiles indicated comparable structural stability between apo and holo systems (Figure S3, Figure S4), suggesting that substrate binding does not compromise overall protein integrity. Binding free energy calculations using the MM/PBSA approach across three independent simulations yielded consistent ΔG_bind values (−27.1, −20.7, and −23.0 kcal/mol), with an average of −23.6 ± 3.3 kcal/mol, supporting a thermodynamically favorable interaction (Table S1). It must be noted that these values should be interpreted qualitatively, as MM/PBSA estimates of absolute binding affinities carry known systematic uncertainties. Energy decomposition indicated that binding was primarily driven by van der Waals interactions, highlighting the importance of hydrophobic contacts between PET-KR1 and the aromatic backbone of the substrate. Electrostatic contributions were smaller, while solvation effects opposed binding, consistent with partial desolvation upon complex formation. Residue-level interaction analysis across MD trajectories identified a set of putative PET-binding residues located around the catalytic Ser132, forming a defined substrate-binding pocket (Figure 8). Residues exhibiting high average interaction occupancy (>80%) across all three replicas were classified as strong interactors, while those observed interacting in only two replicas were considered transient contributors. Key residues consistently involved in PET binding included Phe59, Ile180, Phe211, Thr60, and Met133. Structural comparison with well-characterized PETases further supports the functional relevance of these interactions. Specifically, Phe59 of PET-KR1 aligns with Tyr87 of IsPETase and Tyr95 of LCC, residues previously implicated in π–π interactions with the terephthalate moiety of PET ^79^. Similarly, Met133 and Ile180 correspond to residues in IsPETase (Met161 and Ile208) and LCC (Met166 and Val212) that have been proposed to contribute to substrate binding through hydrophobic interactions ^80^. Phe211 of PET-KR1 aligns with Phe243 of LCC, which has been suggested to stabilize aromatic rings of PET ^80,81^. Additional residues identified in this study provide further insight into substrate recognition. Thr60, conserved across PET-KR1, LCC, and IsPETase, may contribute to binding through hydrogen bonding with ester or terminal groups of the PET oligomer, although its role has not been previously characterized. Conversely, Trp157, corresponding to Trp185 in IsPETase and Trp190 in LCC, displayed high interaction occupancy but was only present in two of the three replicas, suggesting a more transient role in substrate stabilization. Overall, the identified interaction network is consistent with known PETase binding motifs and highlights the importance of hydrophobic and aromatic interactions in substrate recognition. These findings demonstrate that MD simulations provide valuable insight into the dynamic nature of PET binding, complementing static docking predictions and supporting the observed catalytic activity of PET-KR1.

**Figure 8.**
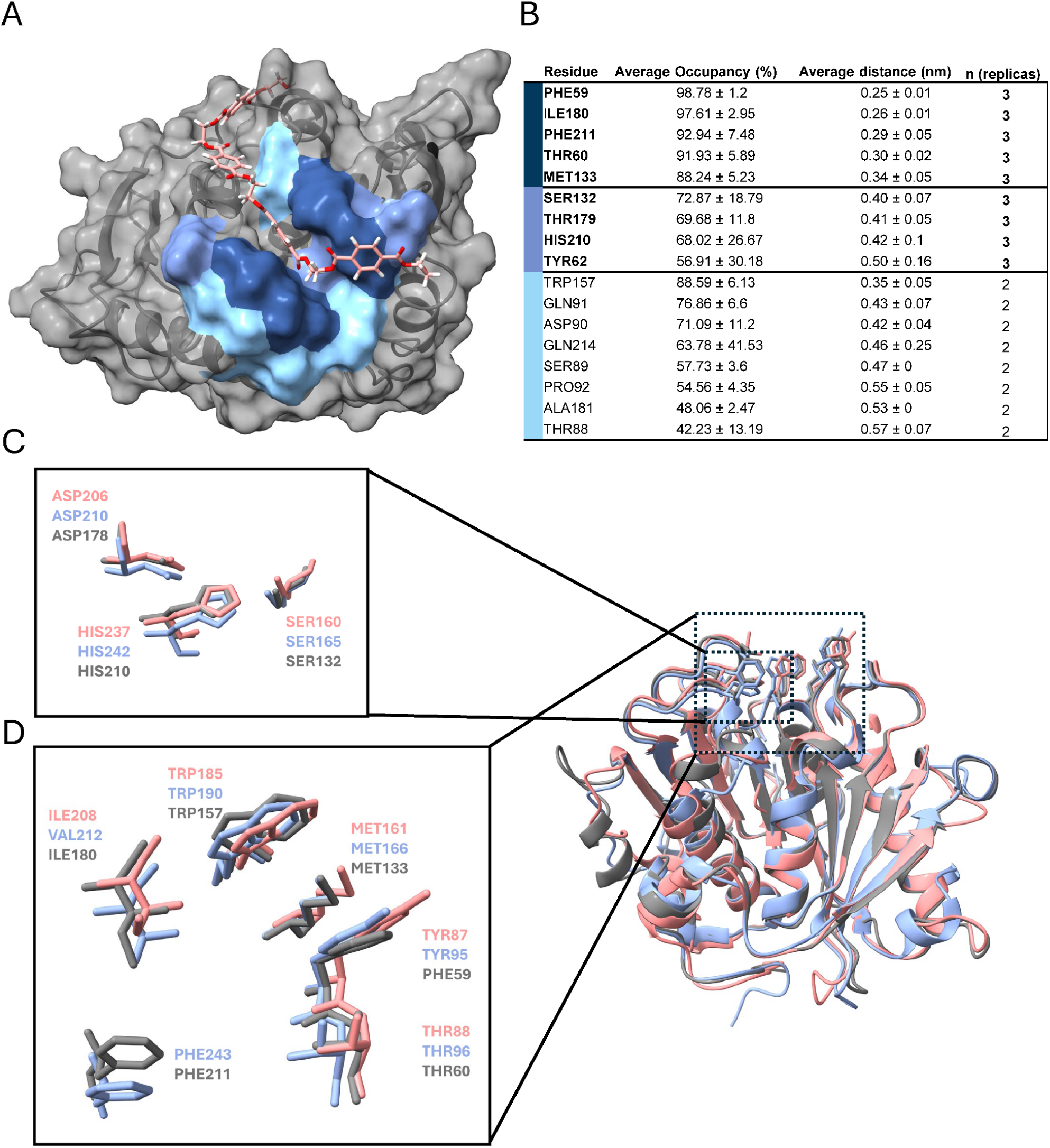
Molecular dynamics analysis of PET binding in PET-KR1 and structural comparison with characterized PETases. **(A)** AlphaFold-predicted structure of PET-KR1 shown as a gray surface with the best-docked 4PET oligomer pose (shown as pink in stick representation) used to initiate the holo simulations. Residues interacting with 4PET across the three MD replicas are mapped onto the structure and colored according to interaction persistence: strong interactors, dark blue; medium interactors, blue; and transient interactors, light blue. **(B)** Residues retained for final analysis were those exhibiting interaction occupancy ≥ 30% in at least two of the three replicas; the table reports their average occupancy and mean residue–ligand distance across the simulations. **(C)** Structural superposition of PET-KR1 (gray) with IsPETase (pink; PDB: 5XJH) and LCC (blue; PDB: 4EB0), highlighting the conserved catalytic triad region. **(D)** Zoomed view of additional 4PET-interacting residues in IsPETase and LCC that overlap with the majority of the interaction hotspots identified by the MD-based analysis of PET-KR1.

### Engineering potential of PET-KR1

Consistent with previous studies showing that metagenome-derived enzymes can provide robust starting scaffolds for protein engineering efforts ^82^, the results obtained in this study indicate that thermostability of PET-KR1 is likely a major limiting factor during PET depolymerization. Although PET-KR1 exhibits a temperature optimum at 65 °C during pNPB hydrolysis, PET depolymerization experiments showed an apparent process optimum at lower temperatures (45–50 °C). This indicates that further stabilization could improve the overall performance under process relevant conditions. Therefore, rational disulfide bond engineering was investigated as a promising stabilizing strategy.

Given the successful application of disulfide engineering in PETases like LCC, Mipa-P and Kubu-P, PET-KR1 was screened for equivalent stabilizing positions ^23,24^. Candidate disulfide bonds were identified by superimposing the AlphaFold model of PET-KR1 with these enzymes and comparing known stabilizing positions with independent predictions from Disulfide by Design 2 and the Maestro web server tools. While DbD2 identifies disulfide-compatible residue pairs based on geometric criteria alone, Maestro evaluates the thermodynamic effect of introducing the bond through energy minimization and ΔΔG calculation, making them complementary tools for candidate selection. Four candidate disulfide pairs were selected for evaluation. Two candidates (R1 and R3) corresponded to positions previously reported to improve thermostability in related PET hydrolases. In addition, R1 and R2 were the only consensus predictions identified by both computational tools, making R2 an attractive candidate despite the absence of literature precedent. Finally, R4 was included as an additional high-ranking stabilization candidate based on its favorable prediction score.

Among these, the single R1 variant of PET-KR1, corresponding to the N206C/S260C cysteine pair, showed a *Tm* of 70 ℃, representing a modest but reproducible increase in apparent melting temperature of 3.5 ℃ (Figure 9 and Figure S2) relative to wild-type PET-KR1 (*Tm* = 66.5 ℃). Notably, this disulfide pair has repeatedly been validated as a stabilizing site, increasing *Tm* by +9.5 °C in LCC (D238C/S283C), +4.5 °C in Mipa-P (A241C/S286C), and +5.0 °C in Kubu-P (A236C/S281C), and was therefore incorporated into the engineered benchmark enzymes LCC-ICCG, Mipa-PM19, and Kubu-PM12 ^23,24^. The smaller Δ*Tm* gain relative to related PETases may reflect differences in the local structural context of the N206/S260 region, which could influence the effective stabilization achievable by this disulfide.

**Figure 9.**
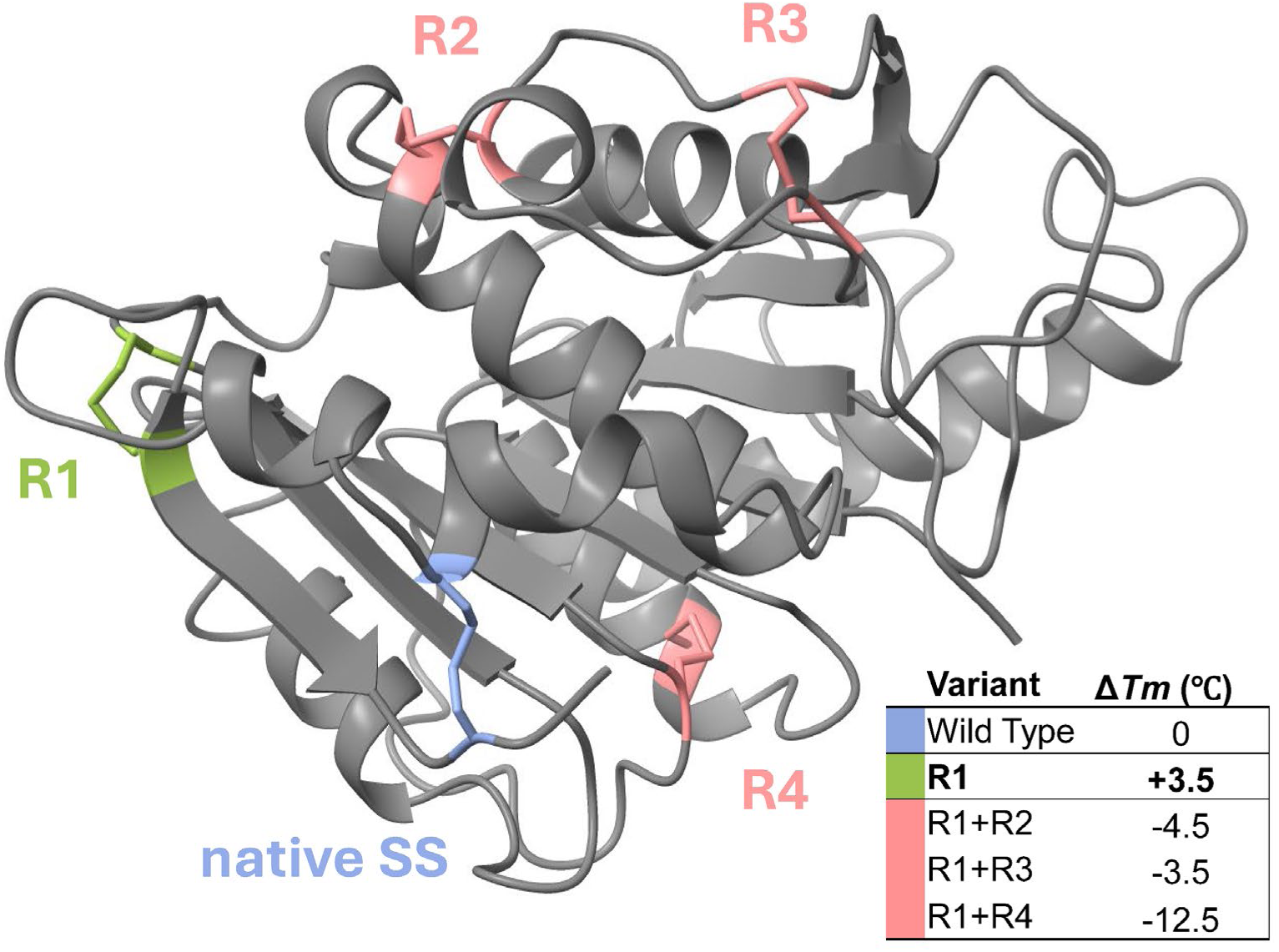
Rational disulfide engineering of PET-KR1. The single native C-terminal disulfide of PET-KR1 is shown in blue, the successfully engineered R1 pair (N206C/S260C) in green, and the remaining candidate disulfides R2 (E15C/P220C), R3 (G6C/P20C), and R4 (A142C/V168C) in pink. The table summarizes the melting temperature shift (Δ*T_m_*) of the variants tested relative to the wild-type enzyme, showing that R1 provided the only stabilizing effect, whereas the combinatorial variants R1+R2, R1+R3, and R1+R4 were destabilizing. The model shown in grey is the AlphaFold-predicted structure of the wild type PET-KR1 while the predicted cysteine substitutions and putative disulfides were modeled in Chimera (v1.19) software, followed by local energy minimization.

By contrast, the combinatorial variants R1+R2, R1+R3, and R1+R4 resulted in lower thermal stability than wild-type PET-KR1, with Δ*T_m_* values of −4.5 °C, −3.5 °C and −12.5 °C thereby offsetting the beneficial effect of the R1 variant. Among these, the promising R3 disulfide pair (G6C/P20C), which corresponds to the stabilizing L49C/S61C pair in Mipa-P and had previously been reported to increase *Tm* by 2.6 °C, did not perform as expected when combined with R1. Overall, our results suggest that PET-KR1 appears to be amenable to protein engineering as supported by the successfully stabilized N206C/S260C variant. Although the stability gains obtained here were modest, the results support further stabilization through additional disulfide engineering, structure-guided stability design and/or directed evolution.

The table summarizes the melting temperature shift (Δ*Tm*) of the variants tested relative to the wild-type enzyme, showing that R1 provided the only stabilizing effect, whereas the combinatorial variants R1+R2, R1+R3, and R1+R4 were destabilizing. The model shown in grey is the AlphaFold-predicted structure of the wild type PET-KR1 while the predicted cysteine substitutions and putative disulfides were modeled in Chimera (v1.19) software, followed by local energy minimization.

## Conclusions

In this study, we developed a targeted metagenomic discovery pipeline and applied it to 277 publicly available plastic-associated metagenomes, shortlisting 21 high-confidence PETase candidates from more than 77 million predicted proteins through successive sequence- and structure-based filtering. Experimental validation identified two novel PET hydrolases, including PET-KR1, a thermostable enzyme (*Tm* = 66.5 °C) that combines a Type I catalytic motif with Type II-like structural features. PET-KR1 depolymerized PET across a broad temperature range with markedly higher productivity on powdered than on film PET substrate, consistent with greater substrate accessibility. While less active overall at higher temperatures (60–65 °C), PET-KR1 generated a product pool more strongly enriched in the terminal monomer TPA. Molecular simulations identified a conserved hydrophobic substrate-binding network around the catalytic serine, and rational engineering of the N206C/S260C disulfide raised thermostability by 3.5 °C, confirming that PET-KR1 is amenable to further optimization.

The wild-type predecessors of current industrial benchmarks were themselves modest performers prior to engineering, underscoring the biotechnological value of expanding the available scaffold repertoire. PET-KR1 represents one such scaffold, while the targeted workflow, built entirely on publicly available tools and open-access data, provides a reproducible strategy for mining plastic-associated metagenomes toward biocatalytic PET recycling.

## Supporting information

Curated ENA Plastic-associated Metagenomes

## Supporting Information

Full multiple sequence alignment of PET-KR1 with characterized PETases; differential scanning fluorimetry (DSF) melt curves of PET-KR1, the engineered PET-KR1 variant (N206C/S260C), and PET-KR2; molecular dynamics analyses (RMSD and RMSF); MM/PBSA binding free-energy calculations; cross-study comparison of PET-KR1 with representative wild-type PETases; and supplementary methods describing the benchmarking analysis (PDF). Curated list of the 277 plastic-associated metagenomic datasets retrieved from the European Nucleotide Archive (ENA), including SRA accession numbers and *ProteoSeeker* screening statistics (CSV).

## Acknowledgements

This research was supported by (i) the Horizon Europe Programme under the “Widening Participation & Spreading Excellence” component (call ERA Chairs “HORIZON-WIDERA-2022-TALENTS-01-01−ERA Chairs”); Project “Boost4Bio”; Grant Agreement No. 101087471, support for K.R., D.B., D.Z., and G.S.; and (ii) the Horizon Europe Programme under the “Widening Participation & Spreading Excellence” component (call Twinning “HORIZON-WIDERA-2021-ACCESS-03-1”); Project “Twin4Promis”; Grant Agreement No. 101079363, support for D.Z. and G.S.; I.A. and D.Z. would like to thank the Cost Action Programme COZYME - COmputationally assisted design of enZYMEs (CA21162) for supporting a research mobility that initiated the collaboration between Luleå University of Technology and the Institute for Bio-innovation, Biomedical Sciences Research Center “Alexander Fleming“.

## Additional Information

Accession codes: The nucleotide sequences of the genes encoding PET-KR1 and PET-KR2 reported in this study have been submitted to the DDBJ/ENA/GenBank databases and are currently under review. Accession numbers will be provided upon release by the database and updated in the final published version of the manuscript.

## Table of Contents (TOC)/Abstract Graphic

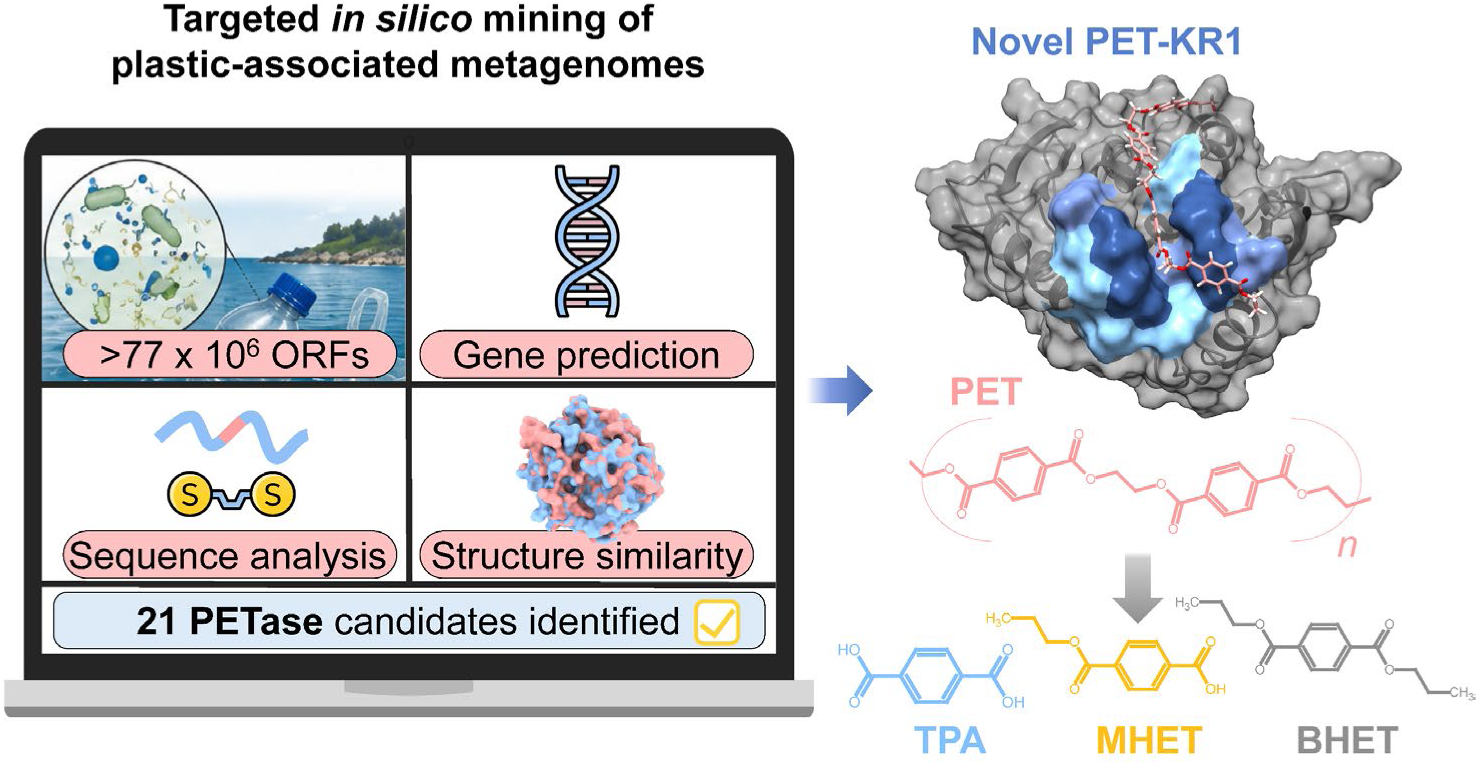

