## Supporting Information for "Targeted mining of plastic-associated metagenomes uncovers a novel thermostable PETase expanding scaffold space for engineering"

### Supplementary materials

#### Figure S1

```
LCC_(Type_I)

LCC_(Type_I)           1  ............MDGVLWRVRTAALMAALLALAAWALVWASPSVEAQSNP...........
PHL-7_(Type_I)         1  ............MA..................................NP...........
Mipa-P                 1  ............M................................................
Kubu-P                 1  ............M................................................
PETcand3               1  ............MA..............................AQADNP...........
PETcand4               1  ............MAG............GGVAAAPAATE.SPAAIQLAANP...........
PETcand5               1  ............MQ.................................TNP...........
PETcand6               1  ............MT...............................TAAQP...........
PETcand7               1  ............MD...................................P...........
IsPETase_(Type_IIb)    1  ............MNF......PRASRLMQAAVLGGLMA.VSAAATAQTNP...........
RgPETase_(Type_IIb)    1  ............MFG........KLPFARASLAVGALL.LSAAAVAQTNP...........
PETcand20              1  .............................................MQTNP...........
PETcand21              1  .............................................MQTNP...........
PE-H_(Type_IIa)        1  ............MPF......NKKSVLA..LCGAGALL.FSMSA.LANNPAPT...DPG.D
PET2_(Type_IIa)        1  ............MPI......TARNTLASLLLASSALL.LSGTA.FAANP.PGGDPDPG..
PETcand15              1  .............................................MQTNP...........
PETcand16              1  ............MG..................................NPDPDPDPDPE.P
PETcand17              1  ............MI..................................TPDPEPDPDPDPP
PETcand18              1  ............M................................................
PETcand19              1  ..............................................MNNPAPT...DPG.D
PET27_(Type_I*)        1  ............MK................KIYAVALF.......FLATS...........
PET30_(Type_I*)        1  ............MR................KLYLFLFL.......TLISP...........
PETcand1               1  ............MQ...............................................
PETcand2               1  ............MQ...............................................
PETcand8               1  ............MQ...............................................
PETcand9               1  ............M...................................NP...........
PETcand10              1  MAALLGTAGAAHAQ...............................................
PETcand11              1  ............MQ...............................................
PETcand12              1  ............MQ...............................................
PETcand13              1  ............MQ...............................................
PETcand14              1  ............M................................................
```

```
                                    η1            β1     η2          β2       ★    β3
LCC_(Type_I)

LCC_(Type_I)          39  ......YQRGPNPTRSALTAD.GPFSVATYTVSRLSVSG..FGGGVIYYPTGT.SLTFGGI
PHL-7_(Type_I)         5  ......YERGPDPTESSIEAVRGPFAVAQTTVSRLQADG..FGGGTIYYPTDTSQGTFGAV
Mipa-P                 2  ......TERGLAPTAANITGD.GSYGVVSATITG..ASG..FGGGVVYYPNAT..ERFPVV
Kubu-P                 2  ......DQVGQAPTAANITGD.GSFATASAPITN..QTG..FGGGTVYYPTAA..GTYPVV
PETcand3              10  ......YERGPAPTEQSIEALRGPFTTSQTSVSSLVVSG..FGGGTIYYPTSTAEGTFGAV
PETcand4              27  ......YERGPAPTERSITAERGPFDIEKIDVPARSGEG..FNQGTIYAPTDLSQGTFGAI
PETcand5               6  ......YQKGPDPTSAALNATSGPFSVATSSVSSL.VIG..FGGGTIYYPTTT..GTYAVL
PETcand6               8  ...GTLVEHGPNPTAQSVQAQRGPYATSRTSASGGLGSG..FNRGTIYYPTTTSEGTFSAV
PETcand7               4  .....AWWRGPAPTEASVSATTGPFATTSTTVSDYSTRG..FGAATIYHPTSTTQGTFGGV
IsPETase_(Type_IIb)   32  ......YARGPNPTAASLEASAGPFTVRSFTVSR..PSG..YGAGTVYYPTNA.GGTVGAI
RgPETase_(Type_IIb)   30  ......YQRGPDPTVSSLEATRGPFSTSSFTVSR..PSG..YGAGTVYYPTNA.GGKVGAI
PETcand20              6  ......YARGPNPTAASLEASTGPFAYRSFTVSR..PSG..YGAGTVYYPTNA.GGTVGAI
PETcand21              6  ......YARGPNPTAASLEASAGPFTVRSFTVRR..PSG..YGAGTVYYPTNA.GGTVGAI
PE-H_(Type_IIa)       36  SGGGSAYQRGPDPSVSFLEADRGQYSVRSSRVSSL.VSG..FGGGTIYYPTGT.TGTMGAV
PET2_(Type_IIa)       39  CQTDCNYQRGPDPTDAYLEAASGPYTVSTIRVSSL.VPG..FGGGTIHYPTNAGGGKMAGI
PETcand15              6  ......YQRGPAPTSSSLAADRGPMAYSQFTVSR..PSG..YRAGTVYYPTNA.GGTVGAI
PETcand16             15  CTSGCDYQRGPDPTTRSLEADRGPYTVRTISVSSFSADG..FGGGTIHYPTNT.TGTMGII
PETcand17             16  STCSNCYQRGPNPTVSALEADSGPYSVRTINVSSW.VSG..FGGGTIHYPVGT.EGTMGAI
PETcand18              2  ......VQIGPNPTKSSLEASRGPFSTSQFTVSR..PSG..YGAGTVYYPTNA.GAKVGAI
PETcand19             12  SGGGSAYQRGPDPSVSFLEADRGQYSVRSSRVSSL.VSG..FGGGTIYYPTGT.TGTMGAV
PET27_(Type_I*)       16  FAPNIIFAQCGDVTIESLTNP.GPYEVATLTEADGLRNGPDYQGATVYYPTNA.TPPFASI
PET30_(Type_I*)       16  ISISIFHAQCTGATVESLTNP.GPYTVATLSEADGVRNGPKYAGSTIYYPTNA.TPPYASI
PETcand1               3  ......YEKGPDPTVSALEASRGPFTIRTSTISRLSASG..FGGGTIYYPASP..GSYGAV
PETcand2               3  ......YEKGPDPTTSALEASRGPFTIRTSTISRLSASG..FGGGTIYYPTAT..GSYGTI
PETcand8               3  ......YQKGPDPTVAALERD.GTFAIRTTTVSRLSASG..FGGGTVYYPTAT..GSYGVV
PETcand9               4  ......YQRGPNPTVAALEASAGPYAVASSKITS..PAG..YGGGTVYYPSGTSDGSFGVV
PETcand10             15  ......YQKGPDPTASALERD.GPFAIRSTSVSSF.VSG..FGGGRLYYPTAT..GTYGAI
PETcand11              3  ......YQKGPDPTASALERN.GPFAIRSTSVSSF.VSG..FGGGRLYYPTAT..GTYGAI
PETcand12              3  ......YQKGPDPTASALERN.GPFAIRSTSVSRF.VSG..FGGGRLYYPTAS..GTYGAI
PETcand13              3  ......FQKGPDPTAAALERD.GPFAIRSTSVSRL.VSG..FGGGRLYYPTAT..GTYGAI
PETcand14              2  ...........DPTTASLEATDGPLSTAGYTVPS..PAG..YGSGTITYPTSS..GSYPGV
```

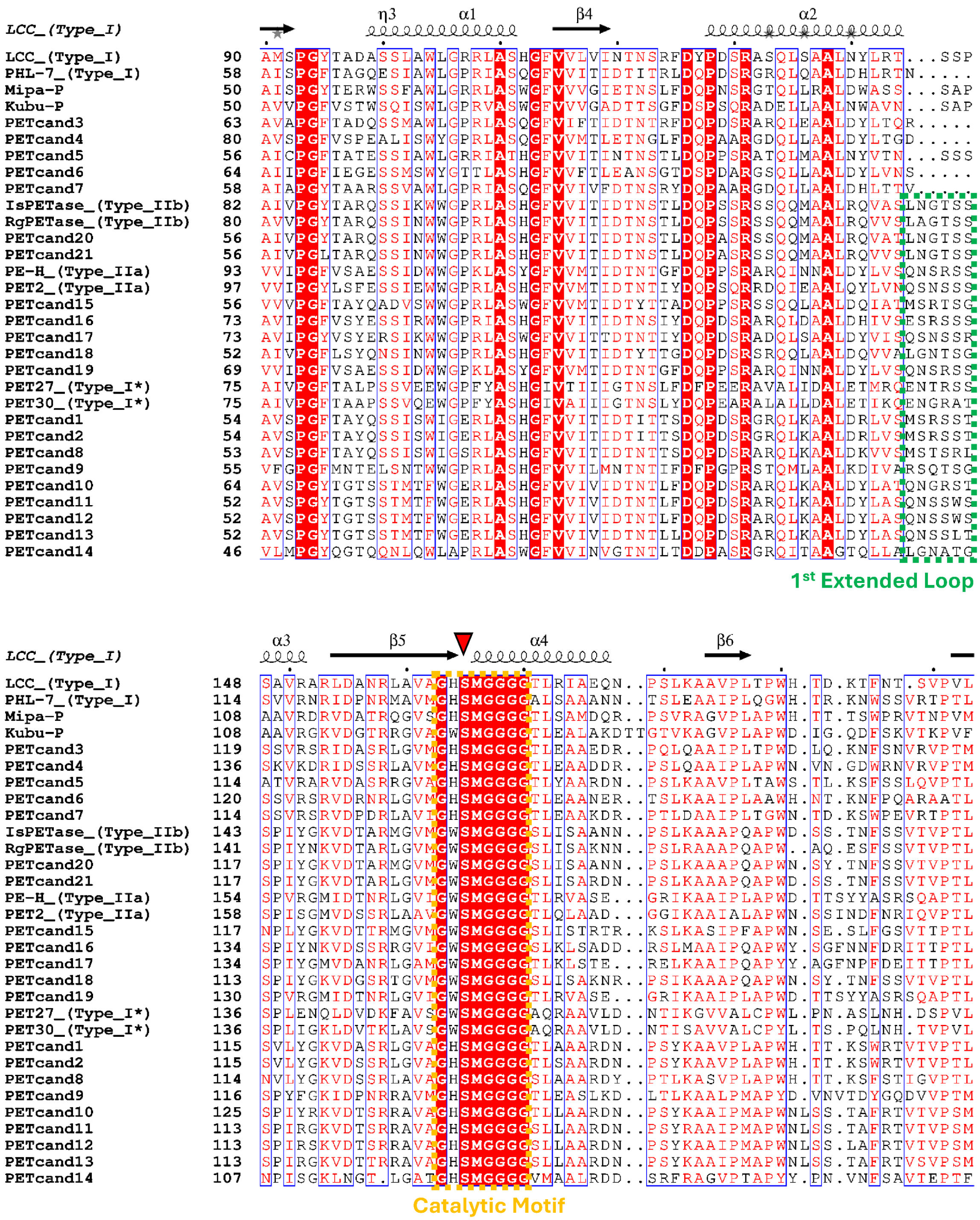


Figure S1 (continued)

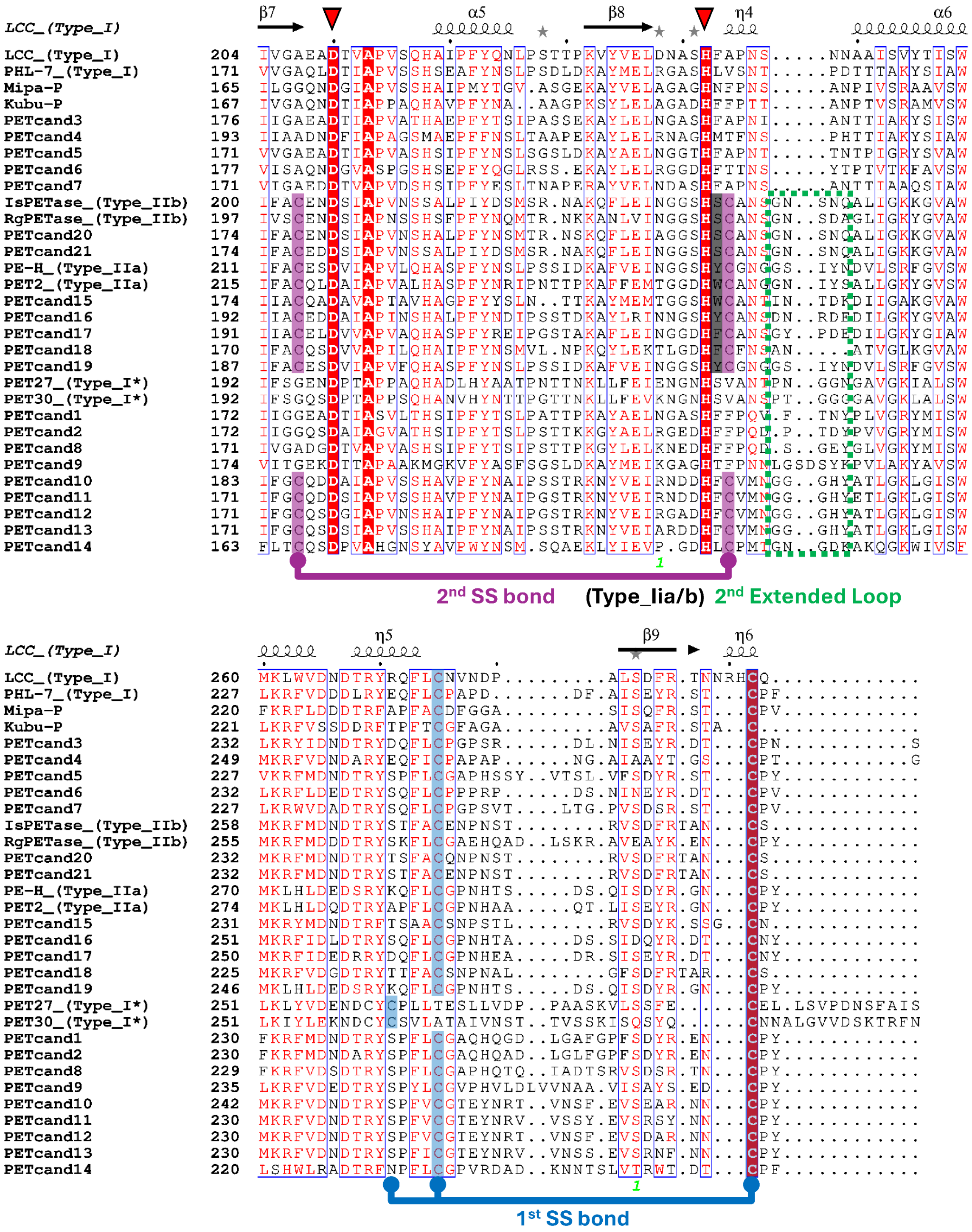


**Figure S1. Full multiple sequence alignment of PET-KR1 with characterized PETases.** The alignment was generated with MAFFT, and conserved sequence features, including the Ser-His-Asp catalytic triad, PETase-defining motifs, C-terminal cysteines, and loop regions associated with substrate binding, are annotated to facilitate comparison across the PETase family. Color coding highlights the same functional features shown in the Figure 3.

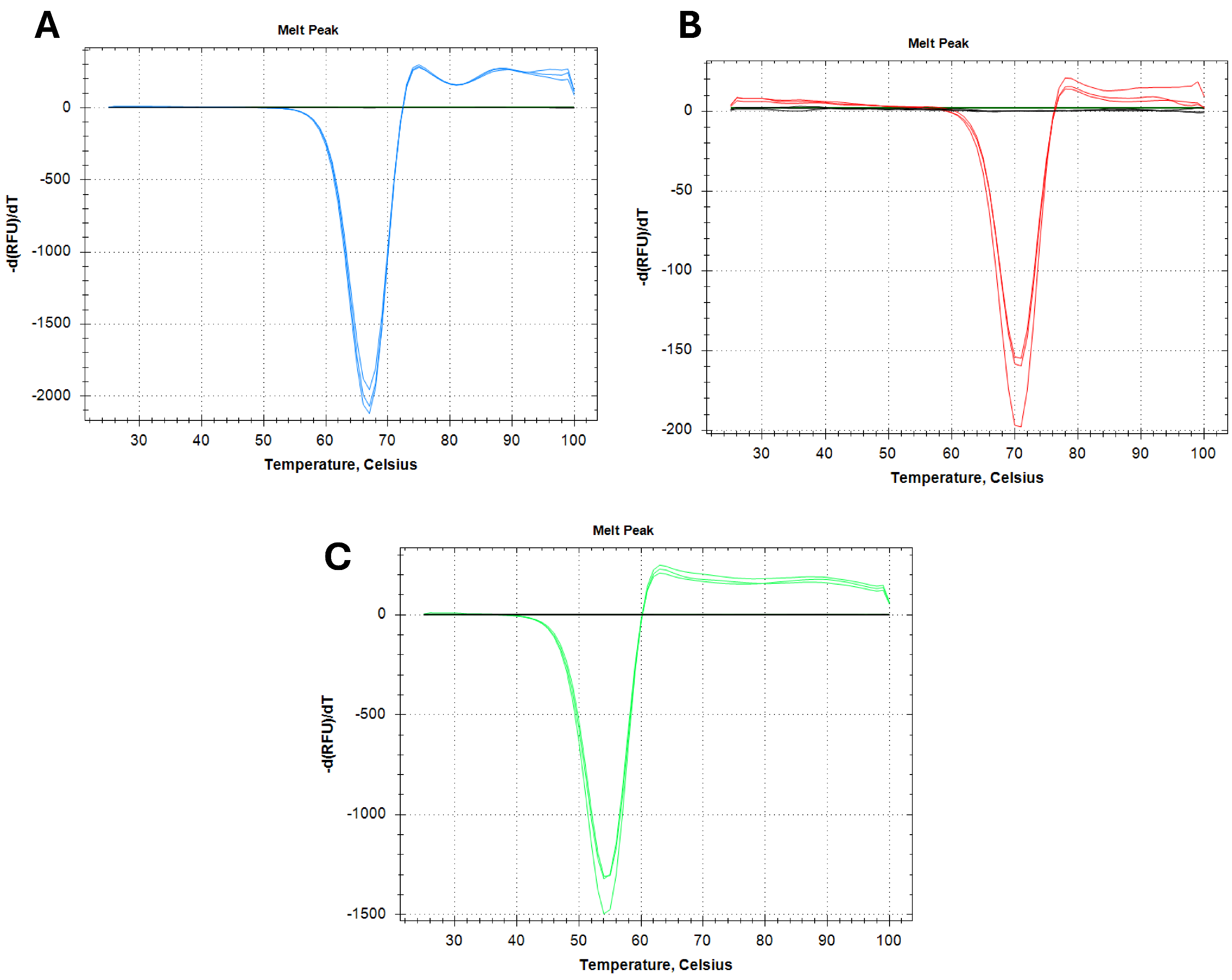


**Figure S2. DSF melt-curve data for novel PETases.** Representative melt curves are shown for **(A)** PET-KR1, **(B)** the successfully engineered N206C/S260C variant of PET-KR1 (R1 PET-KR1), and **(C)** PET-KR2.

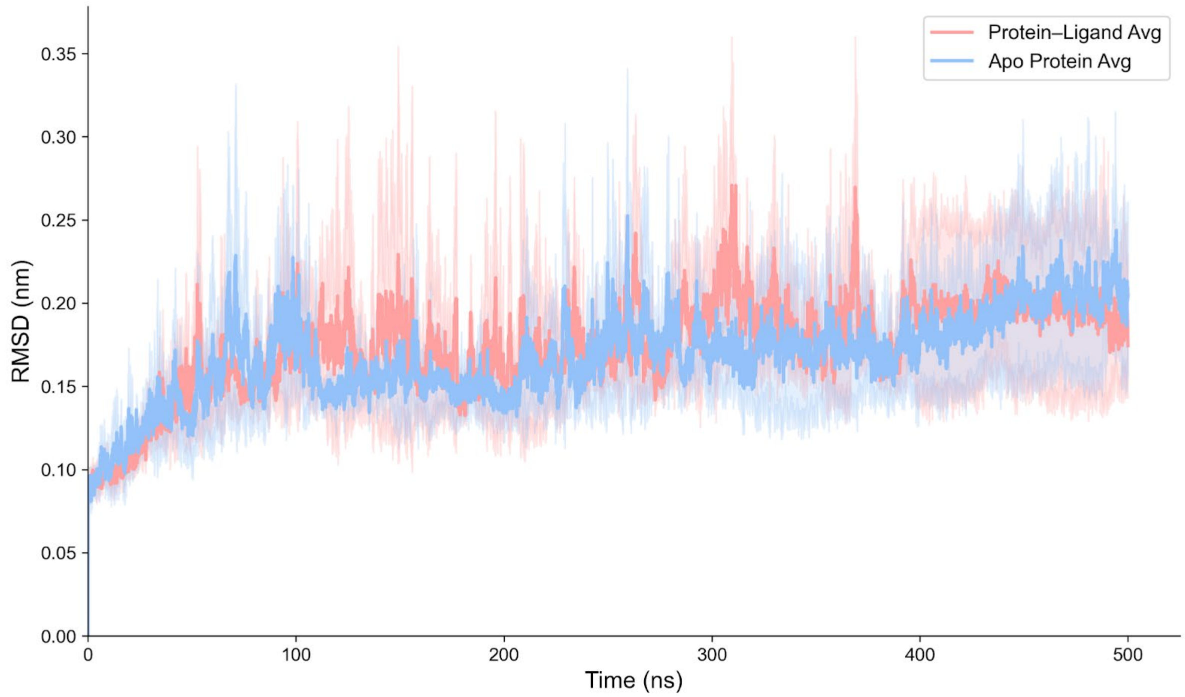


**Figure S3. Root mean square deviation (RMSD).** RMSD profiles of the PET-KR1 protein in the apo (blue) and holo (red) states over the course of molecular dynamics simulations. Shaded regions represent the standard deviation calculated from three independent MD replicas, indicating the variability and convergence of the trajectory.

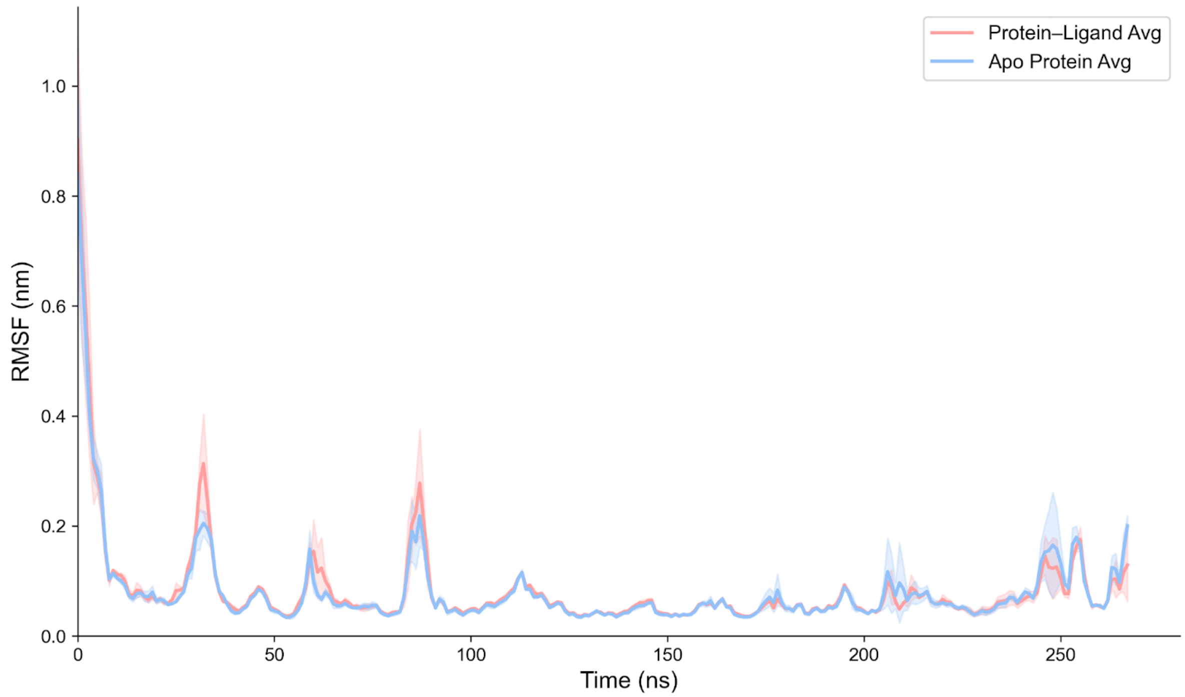


**Figure S4. Root mean square fluctuation (RMSF).** RMSF profiles of PET-KR1 residues in the apo (blue) and holo (red) forms. Shaded areas denote the standard deviation across three independent MD replicas, reflecting residue-level flexibility and the effect of ligand binding on protein dynamics.

**Table S1. Cross-study comparison of PET-KR1 with benchmark wild-type PETases under similar PET-depolymerization conditions.** PET-KR1 is compared with benchmark wild-type PETases: Mipa-P, Kubu-P, CaPETase, IsPETase, and LCC reported by Seo et al. (2025) [1]. Benchmark values were normalized using a midpoint estimate of reported enzyme loading (13–16 µL of 1 mg $mL^{-1}$ enzyme solution). Because substrate loading, PET morphology, and reaction conditions differ between studies, the reported highest specific activities (SA) and specific productivities (SP) are intended for contextual benchmarking rather than strict kinetic equivalence.

| Enzyme | Tm (°C) | Temperature of highest SA (°C) | Highest SA (µmol $h^{-1}$ $mg_{enzyme}^{-1}$) | Temperature of highest SP (°C) | Highest SP (µmol $h^{-1}$ $mg_{enzyme}^{-1}$) |
|---|---|---|---|---|---|
| Mipa-P | 68.6 | 60.0 | 32.0 | 40.0 | 1.6 |
| Kubu-P | 88.8 | 70.0 | 41.3 | 60.0 | 2.4 |
| CaPETase | 64.9 | 50.0 | 8.8 | 40.0 | 1.1 |
| IsPETase | 50.1 | 50.0 | 5.1 | 40.0 | 0.5 |
| LCC | 86.2 | 70.0 | 26.3 | 60.0 | 1.9 |
| **PET-KR1** | **66.5** | **50.0** | **6.5** | **50.0** | **3.6** |

**Table S2. Binding free energy components obtained from MM/PBSA analysis of three independent molecular dynamics simulations**. Reported values represent the mean ± standard deviation across replicas. ΔG_bind corresponds to the total binding free energy (ΔTOTAL), decomposed into van der Waals (ΔVDWAALS), electrostatic (ΔEEL), and solvation (ΔGSOLV) contributions. All energies are given in kcal/mol.

| | MD_Replica1 | MD_Replica2 | MD_Replica3 | **MD_Average** |
|---|---|---|---|---|
| ΔVDWAALS | -40.4 | -31.6 | -34.0 | -35.4 ± 4.5 |
| ΔEEL | -9.2 | -6.9 | -9.3 | -8.4 ± 1.3 |
| ΔGSOLV | 22.4 | 17.8 | 20.2 | 20.1 ± 2.3 |
| **ΔTOTAL** | -27.1 | -20.7 | -23.0 | **-23.6 ± 3.3** |

**Text S1. Comparison of PET-KR1's performance with wild type benchmark PETases**

The highest reported values of initial rate (μM $h^{-1}$ ) and depolymerization extent (μM) at 168 h of reaction were extracted for each benchmark enzyme: Mipa-P, Kubu-P, CaPETase, IsPETase, and LCC from Seo et al. (2025) [1]. The reported enzyme loading (500 nM enzyme; approximately 13–16 μL of 1 mg $mL^{-1}$ enzyme solution per 1 mL reaction) was converted to an effective enzyme concentration using the midpoint value (14.5 μL equivalent, corresponding to ~13 mg $L^{-1}$). Specific activity (μmol $h^{-1}$ $mg_{enzyme}{}^{-1}$) was calculated by dividing reported initial rates by this estimated enzyme loading. Specific productivity (μmol $h^{-1}$ $mg_{enzyme}{}^{-1}$) was calculated by dividing total depolymerization extent at 168 h by reaction time and enzyme loading. This normalization enables cross-study comparison but does not imply strict kinetic equivalence due to differences in substrate loading, PET morphology, and reaction conditions.
